# Quantifying the mechanics and growth of cells and tissues in 3D using high resolution computational models

**DOI:** 10.1101/470559

**Authors:** Paul Van Liedekerke, Johannes Neitsch, Tim Johann, Enrico Warmt, Ismael Gonzales Valverde, Stefan Höhme, Steffen Grosser, Josef Käs, Dirk Drasdo

## Abstract

Mathematical models are increasingly designed to guide experiments in biology, biotechnology, as well as to assist in medical decision making. They are in particular important to understand emergent collective cell behavior. For this purpose, the models, despite still abstractions of reality, need to be quantitative in all aspects relevant for the question of interest. The focus in this paper is to study the regeneration of liver after drug-induced depletion of hepatocytes, in which surviving dividing and migrating hepatocytes must squeeze through a blood vessel network to fill the emerged lesions. Here, the cells’ response to mechanical stress might significantly impact on the regeneration process. We present a 3D high-resolution cell-based model integrating information from measurements in order to obtain a refined quantitative understanding of the cell-biomechanical impact on the closure of drug-induced lesions in liver. Our model represents each cell individually, constructed as a physically scalable network of viscoelastic elements, capable of mimicking realistic cell deformation and supplying information at subcellular scales. The cells have the capability to migrate, grow and divide, and infer the nature of their mechanical elements and their parameters from comparisons with optical stretcher experiments. Due to triangulation of the cell surface, interactions of cells with arbitrarily shaped (triangulated) structures such as blood vessels can be captured naturally. Comparing our simulations with those of so-called center-based models, in which cells have a rigid shape and forces are exerted between cell centers, we find that the migration forces a cell needs to exert on its environment to close a tissue lesion, is much smaller than predicted by center-based models. This effect is expected to be even more present in chronic liver disease, where tissue stiffens and excess collagen narrows pores for cells to squeeze through.

## 1 Introduction

Driven by the insight that multi-cellular organization cannot be explained by the orchestration of chemical processes at the molecular level alone and flanked by recent development of methods in imaging and probing of physical forces at small scales, the role of mechanics in the interplay of cell and multi-cellular dynamics is moving into the main focus of biological research [1]. Cells respond on mechanical stress passively and actively, hence an understanding of growth and division processes is not possible without properly taking into account the mechanical components underlying these processes. Mathematical models are being established as an additional cornerstone to provide information to clinicians entering in their decisions [2, 3]. In particular effects based on nonlinear dynamics need mathematical modeling. This requires high reliability of models and quantitative simulations.

An clinical relevant example is the regeneration of liver after drug induced toxic damage after paracetamol (acetaminophen, APAP or Tetrachloride, CCl4) overdose. These drugs generate a characteristic central necrotic hepatocyte-depleted lesion in each liver lobule, which is the smallest repetitive functional and anatomical unit of liver. Hoehme et al. (2010) [4] used confocal laser scanning micrographs to set up a realistic spatial-temporal model of a liver lobule. In their model hepatocytes were represented as individual units (agents) parameterized by biophysical and biological quantities and able to move as a consequence of forces on the cell and the cells’ own micro-motility. The cells were approximated as spheres, while the forces between them are simulated as forces between the cell centers, which is why the models are often termed “center-based model” (CBM). Center-based models have proven useful to mimic tissue organization processes for example in-vitro and in early development (see e.g. [5–8]) and have been shown to provide a good framework for multi-scale simulations in tumor development (e.g. [9, 10]). For liver, the CBM predicted that active uniform micro-motility forces would not suffice to close the characteristic necrotic tissue lesions generated in the center of each liver lobule but further mechanisms as directed migration and oriented cell division along closest micro-vessels (named hepatocyte-sinusoid alignment, HSA) would be necessary to explain the observed regeneration scenarios. HSA could subsequently be experimentally validated. To obtain these results, extensive simulated sensitivity analyses had been performed varying each parameter of the model within its physiological range [11]. Hence, in order to arrive at such a conclusion, the model must for a given set of parameters be able to realistically and quantitatively predict the outcome of the regeneration process. The model can be viewed as a mapping from a set of parameters to a set of typically macroscopic observables such as the size of the necrotic lesion, the cell density in the lobule etc.

A major drawback of center-based models is that they are based on the calculation of pairwise forces (usually Hertz force, Johnson-Kendall-Roberts force or related) between cell centers, which fails in dense cell aggregates under compression where the interaction force of a cell with one neighbor impacts on its interaction force with another neighbor. Such situation occurs during liver regeneration after APAP or CCl4 intoxication where many cells enter the cell cycle almost at the same time close the drug-induced lesion. It also occurs in the interior of growing multicellular spheroids. As a consequence of considering only pair-wise forces in absence of a notion of cell volume, incompressible cells characterized by a Poisson ratio of *ν* = 0.5 might in a simulation become over-compressed leading to unrealistic multicellular arrangements and thereby to false predictions. Corrections have been proposed to circumvent this shortcoming, but so far a fully consistent approach for center-based models has been been out of reach as this requires to consistently relate cell-cell interaction forces and cell shape. In center-based models, cell shape can only be estimated for very small cell deformations. Approximating cell shape by Voronoi-tesselation [12] permits to calculate a cell volume, but the interaction forces are not necessarily consistent with the cell-cell contact areas resulting from this tesselation [13].

The shortcomings of the CBM call for a model calls for a cell type that consistently relate cell strain and stresses in cells within multicellular assemblies such as in a liver lobule. A large category of models called lattice-free, force-based “Deformable Cell Models”(DCMs) has been developed to meet these needs [14–17]. Their lattice based counterparts, called Cellular Potts Models (CPM), are popular in biomedical modeling, partially because of their straightforward implementation. The dynamics in CPM is principally stochastic in nature and is based on the minimization of energy functionals (Hamiltonian) and Monte-Carlo sampling over a vast number of lattice sites (e.g. [18–20]). Force-based methods use equations of motion, which facilitates presenting both, stochastic and deterministic components. Early work in this field has treated multicellular spheroids, various cellular patterns in developing ductal carcinoma in situ, invasive tumors as well as normal development of epithelial ductal monolayers and their various mutants [14, 15, 21, 22]. Deformable models have also been developed to study the dynamics of erythrocytes in blood flow [23–26], cellular rheology [27] or even impact of tissue [28, 29]. Recent approaches focus more and more on the explicit representation of subcellular details such as a nucleus, and cytoskeleton [16, 30]. Other DCM classes, such as the Vertex Model, focus intrinsically more on epithelial sheets [31].

In this paper we present a high resolution deformable cell model in three dimensions parameterized by physical and bio-kinetic parameters, allowing to simulate cell growth in tissues to deepen our understanding in the mechanisms of tissue regeneration and quantify the relations between cell growth, mechanical variables and tissue architecture. Our cell model builds upon earlier work by [17, 32], whereby the cell surface is triangulated and the nodes are connected by viscoelastic elements, representing the cell membrane and actin cortical cytoskeleton and homogeneous cytoplasm. In the work of Odenthal et al. (2013) [17] it was shown that this DCM could quantitatively mimic the adhesion dynamics of red blood cells on a surface, yet cell growth and division were not envisaged. Here, we enrich the model with those features, enabling us to model various multi-cellular systems. Division of individual, isolated cells displaying shape have been modelled by a number of authors (e.g. [16, 33, 34]), whereby in ref. [16, 35] the mechanical processes leading to cytokinesis have been explictly mimicked. Most of these models are two dimensional. Growth and division in three dimensional triangulated cells turns out to be challenging with regard to both the algorithms and the computational time. Simulation of growth is mimicked by adapting the cells’ viscoelastic elements to the new intrinsic size. Cytokinesis, during which a cell splits into two separated daughter cells, completes the mitosis phase. Mitosis and cytokinesis together take about 1*h* compared to the duration of the cell cycle ∼ 24 *hrs* in most mammalian cells, hence are very short. For this reason, we mimic the split of one into two cells in one step however ensuring that the daughter cells precisely fill the space of the mother cell.

By direct comparison with optical stretcher experiments [36], which cause cell deformation as a response of a externally applied stress, we determine the type of the cell’s viscoelastic elements and the magnitude of its parameters. In addition to the basic cortical triangulated model, we have created the possibility for each cell to mimic an internal cytoskeleton by connecting the cell cortex and the cell nucleus by viscoelastic elements. However, as we have found that the deformation of cells can quantitatively be captured even without explicit representation of cell nuclei, we perform the simulations in this paper without explicit representation of the cell nucleus and elements linking cell nucleus and cortex.

The model allows to simulate simple structures as spheroids and monolayers, in which cells interact basically only with each other or with a large rigid plane. In liver architecture, relevant for our final application example, cells interact with other cells, but also with a complex network of micro-vessels (named “sinusoids”). Therefore our design is such that each cell can interact with an arbitrarily shaped object that is either triangulated or represented by a smooth mathematical surface. This also facilitates performing hybrid tissue simulations where DCM cells can interact with center-based cells. Generally, hybrid simulations are useful if simulations of an entire tissue need to be performed in a reasonable time [37]. Hybrid modeling combining DCM and center-based model, conceptually similar to the hybrid strategy proposed in [38], enables us to simulated part of the system as higher spatial resolution and thereby “zoom” into spatial substructures of interest. To ensure that the center-based model behaves “on average” as the DCM, which is a prioi not the case due to the shortcomings of the CBM approach at high cell densities discussed above, we here propose a simple correction scheme in which the interaction forces of the CBM are calibrated with simulations with the DCM. In this way the DCM can be used to verify the systems behavior of the CBM for small cell populations, while the CBM can be used to simulate large cell populations. We demonstrate this by direct comparison of a CBM and a DCM in the same liver lobule.

This paper is structured as follows. The technical details of the DCM and the CBM (and force calibration) are explained in section 2. Following, we first study the single cell dynamics of the DCM. Each model parameter, biomechanical and biokinetic, can be directly associated with a physical property i.e. can either be directly measured or be calibrated by comparison to single cell or multi-cellular experiments. This makes it possible to identify physiological parameter ranges. We here compare directly optical stretcher real to *in silico* (with the DCM) experiments (section 3.1.1) to identify the nature of visco-elastic elements in the DCM and their parameters. Next, we consider classical in-silico experiments of two adhering cells being mechanically separated to identify the model parameters for cell-cell adhesion. We verify whether the contact forces and stress distribution on the cell surface predicted by our model are physically plausible (section 3.1.2). These experiments can generally serve as means to calibrate the parameters of a single cell accurately.

Secondly, we consider the growth and division behavior of the cells in the classical settings of growing monolayer and multicellular spheroids, which have been studied in numerous experiments and modeling works (section 3.2).

Finally, we perform simulations of regeneration dynamics after intoxication of liver with CCl4/APAP using the DCM and compare simulation results to both experimental data and simulation results with the CBM similar as in ref. [4] described above. We analyze the results and basic differences in terms of dynamics and tissue architecture in section 3.3. The cell shapes obtained by simulations with the DCM can in principle readily be compared with high resolution confocal microscopy images. Together with developments in tissue clearing [39], might open up the possibility to infer the stresses on the cells in full 3D volume reconstructions from laser scanning confocal micrographs of the cell shapes. Alternatively, the elastic properties of emergent tissues simulated with the model can be compared to elastographic images [40].

## 2 Mathematical models

In agent-based models of multi-cellular assemblies every cell (= agent) is represented as an individual separated, usually geometrical, object that is able to move, grow, divide, and die. The cell can interact with other cells as well as other objects in its environment. As such, emerging effects of these many interactions can be studied. We explain the two types of single cell-based models used in this work: first the deformable cell model (DCM), then we recapitulate center-based model (CBM).

### 2.1 Deformable Cell Model (DCM)

In our version of a DCM, the cell surface is triangulated with viscoelastic elements along each edge of each triangle. This creates a deformable structure with many degrees of freedom for cell surface deformation [17, 28, 32]. Throughout this paper, we do not represent the cell organelles separately but by a homogeneous isotropic viscoelastic material, see Fig. 1A, despite our model in principle permits the explicit representation of organelles e.g. by triangulating them in the same way as the cell surface and connecting the structures by viscoelastic elements (Fig. 1B).

**Fig 1.**
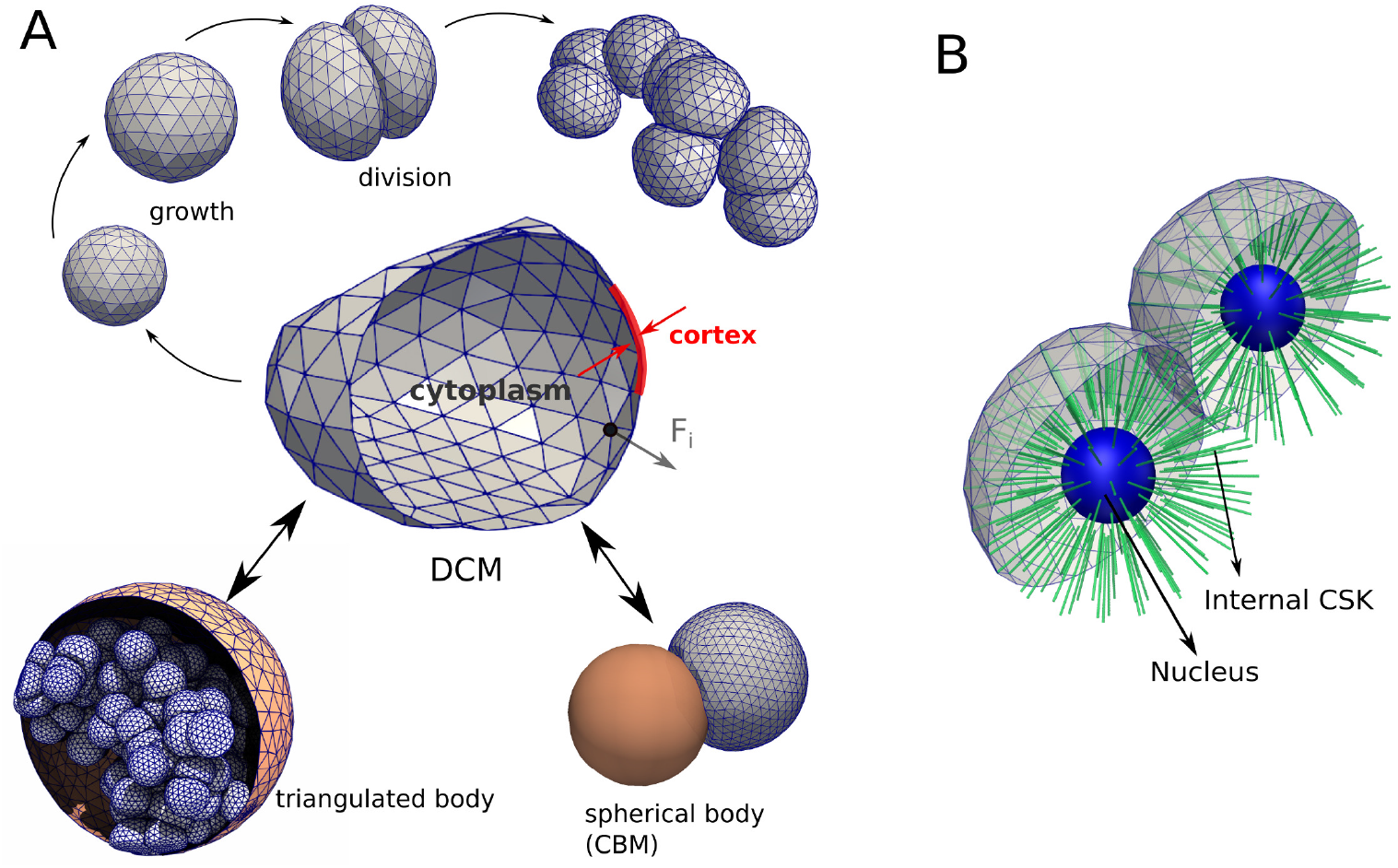
(A) The force-based Deformable Cell Model (DCM) and its basic components and functionality used in this work. A cell is represented by a viscoelastic triangulated shell (cortex) containing a compressible cytoplasm (center). The cells can grow until they split into two new cells, eventually creating a clump of adhering cells (top). Each nodal point of the cell move according to an equation of motion in response to a force **F**_*i*_. The cells can interact with rigid triangulated bodies (such as here a capsule encapsulating them) or simple geometric bodies such as a center-based model (bottom). (B) Same model showing prototype of the model where a nucleus and internal cytoskeleton is included explicitly.

We found that the homogenized approximation as the explicit representation of cell organelles was both not needed and computationally too costly for the questions studied in this paper. The viscoelastic elements and parameters of the cell surface are calibrated such that they simultaneously accounted for the mechanical response of the cell membrane and cell cortex, in particular the cortical cytoskeleton (CSK) i.e., forces that represent in-plane and bending elasticity of the cortex and the plasma membrane. Moreover, we account for a force summarizing contributions from the cytoplasm and the nucleus in response to cell compression. Cell-cell interactions induce external forces, which may be repulsive or adhesive, or both.

Cells in our model can grow and divide when their volume has doubled. When a cell divides, two new cells are created that fit in the envelop of the mother cell and all aforementioned forces are automatically invoked on the daughter cells (see Fig. 3). Apoptosis can be modeled as well. The details of the model components are explained in the following sections.

**Fig 2.**
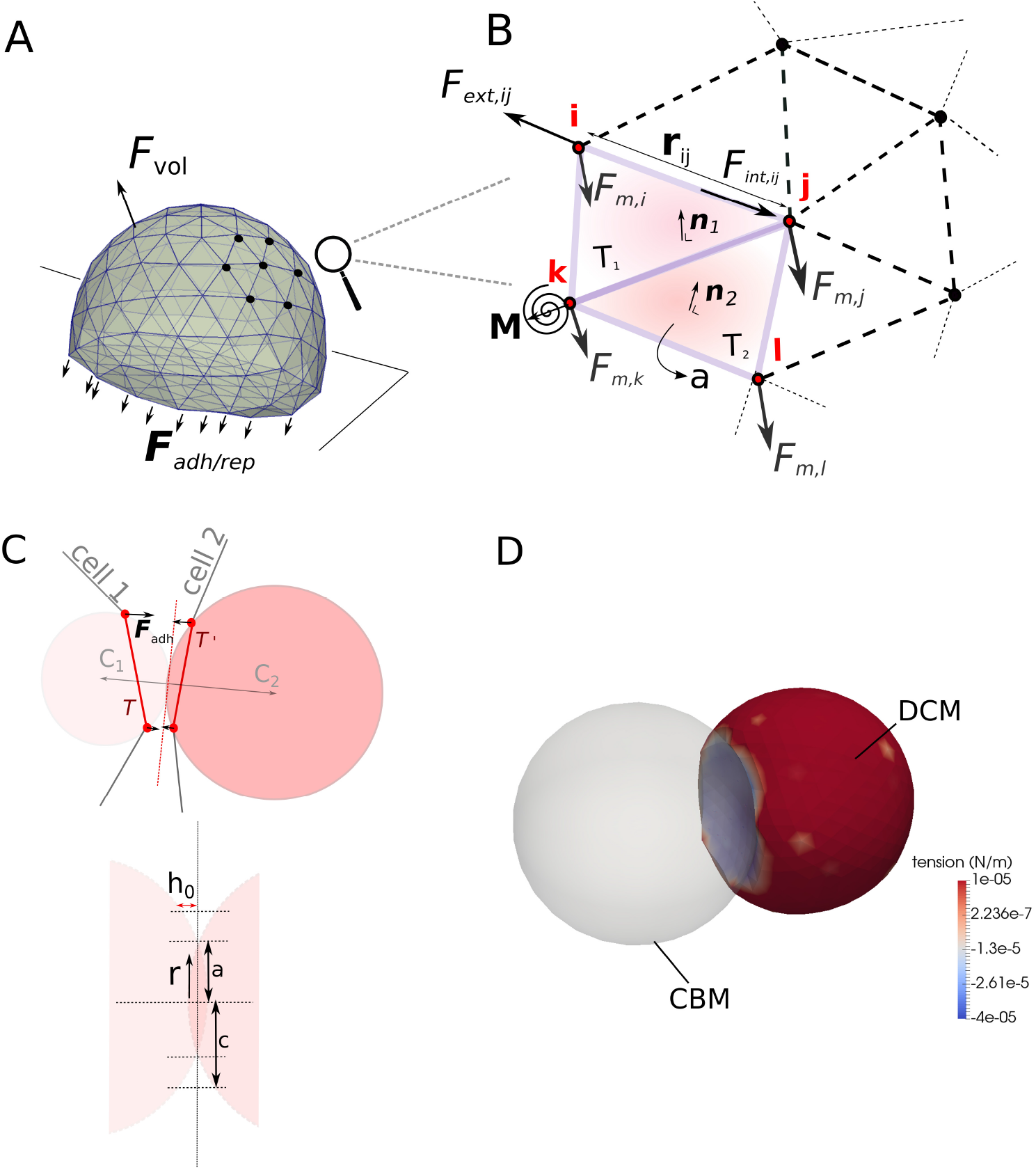
(A-B) Cartoon of adhering cell and detail of the force elements acting in the cell surface triangulation. (C) 2D representation of a triangle pair (T,T’) belonging to different cells to compute the cell-cell interaction force **F**_*adh*_. The 3 nodes of a triangle are here projected as two nodes, whereas the circumscribing spheres are represented as circles. (D) Simulation snapshot of adhesion between a CBM and a Deformable Cell Model (DCM). The colorbar is according to membrane tension (see Eq. 8).

**Fig 3.**
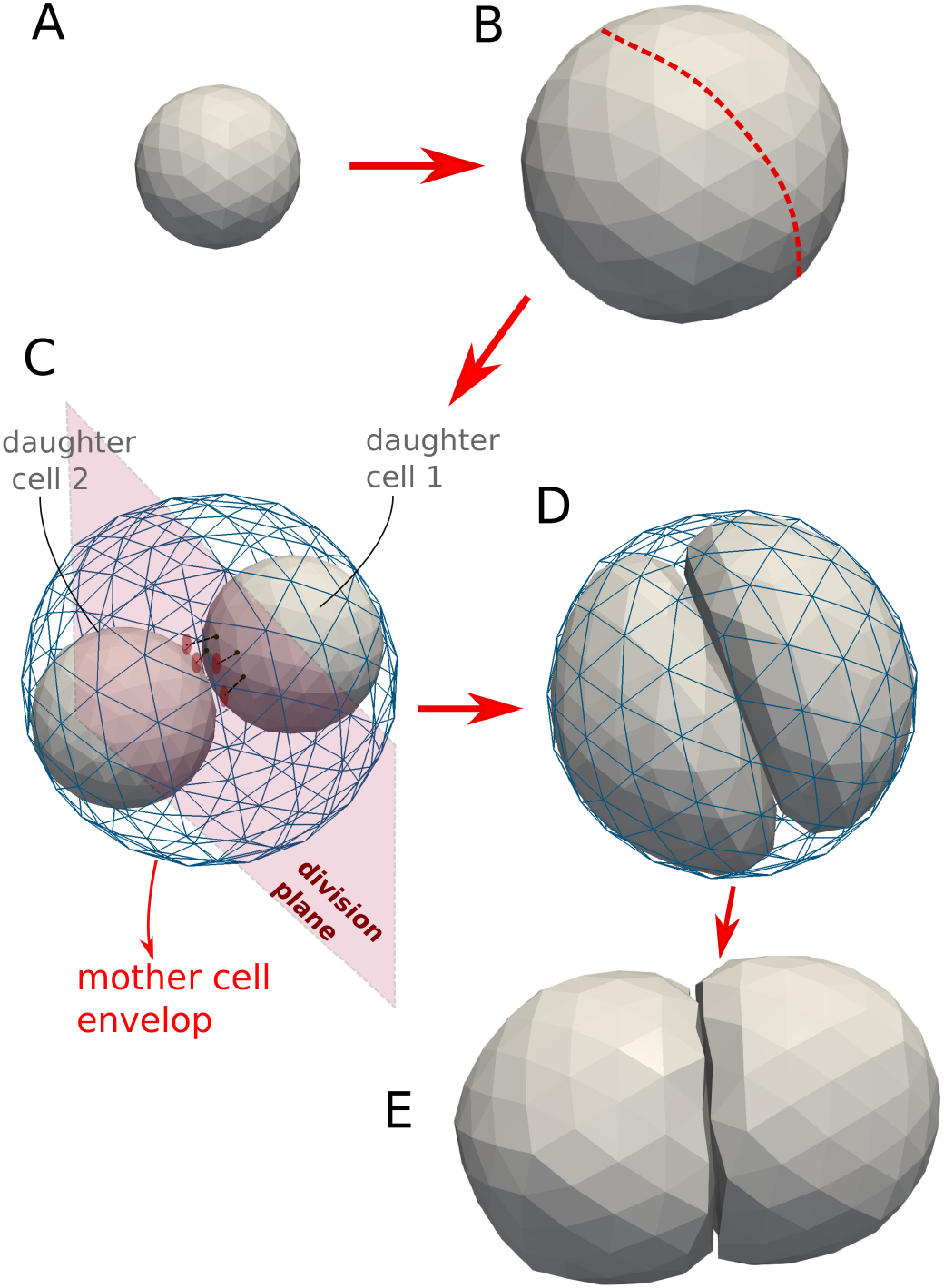
Model mechanisms of the cell cycle of deformable cells: (A-B) Cell growth (double volume by increasing the radius). (B) Choose division direction randomly (assign nodes to one side). (C) Add two default cells as daughter cells to center of masses. Project nodes of the daughter cells to surface of division plane if they intersect it. A sub-simulation is run to come rapidly to situation D. (D) After the sub-simulation, mother envelope is removed. (E) Division stage finished: two mechanically relaxed daughter cells.

#### 2.1.1 Forces and equations of motion

Movement and deformation of a cell can be calculated from a force balance summarized in the following equation of motion for each node *i*:

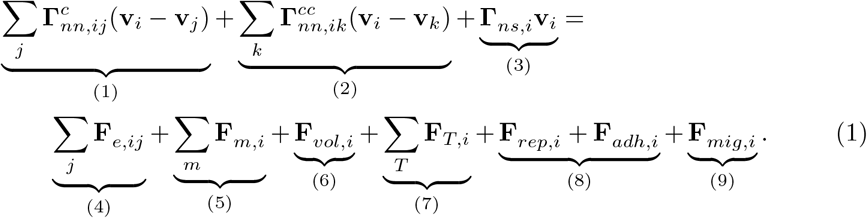

The terms denote (1) node-node friction of nodes belonging to the same cell, (2) node-node friction of nodes belonging to different cells (alternatively, the first two terms as they have the same form could be casted into one term keeping in mind that friction coefficients among nodes of the same cell and of a cell with another cell may differ), (3) friction between nodes of the cell and extracellular matrix (ECM) or liquid, (4) CSK in-plane elasticity, (5) CSK bending, (6) a volume force term penalizing deviations from the cells’ intrinsic volume, (7) a membrane area conservation force, (8) a contact force consisting of a repulsive and adhesive part with a substrate or other cells, and (9) an active migration force. Inertia terms have been neglected as the Reynolds numbers of the medium circumventing the cells are very small [41]; this approximation is common for cell movement (see e.g. [13, 17]). More specifically, the matrices **Γ**_*nn*_ and **Γ**_*ns*_ represent node-node friction and node substrate (ECM) friction, respectively. **v**_*i*_ denotes the velocity of node *i*. The first and the 2nd term on the rhs. represent the CSK in-plane nodal elastic forces **F**_*e,ij*_ and the bending force **F**_*m,i*_. Cell contractility can be included in the model as an extra active elastic force term, but here we assume the cell is in a contractile neutral state. The third term on the rhs. is a volume force **F**_*vol,i*_ and controls the cell compressibility. We assume that cells are compressible on longer timescales controlled by in-and outflow of water. As water transport volume flow rates are small, on short time scales cell volume can be only slightly compressed by compression of the elastic structures inside the cell such as the cytoskeleton, hence the cell exhibits a near incompressible behavior. On longer timescales, the cell response may become more complex due to intracellular adaptations [42]. The force **F**_*T,i*_ accounts for resistance of isotropic expansion of the cell membrane. The two terms (**F**_*adh,i*_, **F**_*rep,i*_) account for potential adhesion and repulsion forces on a local surface node, exerted by an external object such as other triangulated cells or rigid structures (see Fig. 1A). **F**_*mig,i*_ describes the migration forces acting on each node to result in a global movement of the cell. We now give more detail on how these forces and friction components can be calculated.

*Friction terms:* The matrices 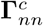, 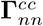 and **Γ**_*ns*_ in Eq. 1 represent node-node friction and node substrate friction tensors, respectively. Individual friction coefficients for two nodes are denoted as *γ_k_* where *k* can refers to the nature of the contact (i.e., nodes on the same cell; nodes of two different cells; friction of a node with extracellular matrix; friction of a node with a liquid). Furthermore, we distinguish between friction in parallel (*γ_||_*) and normal (*γ*_⊥_) direction to the relative motion. If the parallel and normal coefficients are not equal (anisotropy), the friction tensor becomes

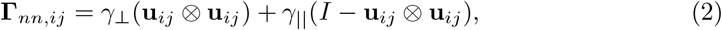

with **u**_*ij*_ = (**r**_*j*_ − **r**_*i*_)/||**r**_*j*_ − **r**_*i*_||, where *r_i_*, *r_j_* denote the position of the nodes of a cell. *I* is the 3 *×* 3 identity matrix, while ⊗ denotes the dyadic product. The nodal cell-substrate friction Γ_*ns*_ can represent viscous resistance with liquid, ECM, capillaries or membranes. If the cell is spherical and the medium is isotropic, then Γ_*ns*_ is a diagonal matrix^1^. It is reasonable to split the friction coefficients -that have unit *N s/m*- into a product of a friction coefficient per surface area (unit *N s/m*^3^) and the shared surface area associated with the interaction of the two nodes (unit *m*^2^). The nodal areas are calculated in the contact model (see further below). In case of friction of a node *i* with an external medium this is the surface area is the Voronoi area of that node (see [28]).

*Cystoskeleton in-plane elasticity - term (4) in Eq. 1:* The CSK in-plane elasticity is controlled by the viscoelastic elements. The elastic forces in the viscoelastic network of a cell cortex are here represented by linear springs with spring stiffness *k_s_*, while the friction coefficients are denoted as *γ_int_* and might be represented by dashpots. Complex viscoelastic elements can be constructed by combining several springs and dashpots or using nonlinear springs [17]. We here do not assume an active contractile state of the cell.

We briefly summarize the components necessary for the Kevin-Voigt Element (Fig. 5D). The combination in which the spring and friction terms are schematically positioned in parallel to each other reflects the Kelvin-Voigt Element (KVE), representing a solid like behavior so that the elements after release of an external force relaxes back to its original length. Consider for example the internal force *F_int_* originating from the Kelvin-Voigt viscoelastic element between node nodes *i* and *j* (for simplicity *γ*_||_ = *γ*_⊥_ = *γ*).

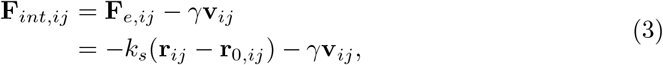

where **r**_0,*ij*_ and **r**_*ij*_ are the initial and actual distance vectors, and **v**_*ij*_ = **v**_*j*_ − **v**_*i*_ is the relative velocity. The force balance equation with external forces **F**_*ext*_ demands that **F**_*ext*_ + **F**_*int*_ = **0**, hence:

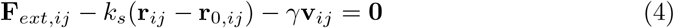

Contrary, a Maxwell element (ME) simulates a fluid like extension of the cortex. In the ME the friction element and spring element are schematically positioned in series (see Fig. 5D), which is why an external force leading to an extension of the element is damped but removal of the force does not lead to relaxation of the element back to its original length characteristic for a fluid behavior. The equation of the internal force in the element **F**_*int*_ (assuming isotropic constants) involves a differential and reads

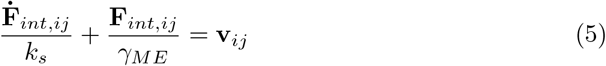

This equation contains a derivative of the force, which can be approximated by **Ḟ**_*int*_ = (**F**_*int*_(*t*) – **F**_*int*_(*t –* ∆*t*))*/*∆*t*. From the force balance between external and internal forces (Eq. 4) we find now

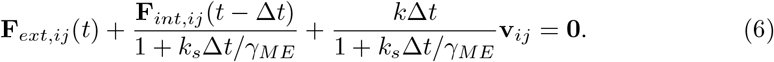

This equation thus involves the evaluation of the internal force on the previous timestep.

The linear spring constant for a sixfold symmetric triangulated lattice can be related approximately to the cortex Young modulus *E_cor_* with thickness *h_cor_* by [43]

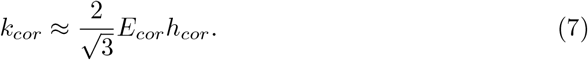

Furthermore, the total elastic in-plane forces can be related to a local in-plane stress using the formula [17]:

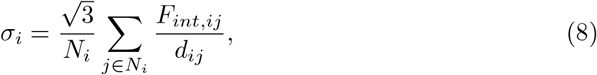

where *N_i_* is the coordination number of node *i*, *F_int,ij_* is the total elastic force between *i* and *j*, and *d_ij_* is their distance.

*The membrane area conservation force - term (6) in Eq. 1:* The force component on the element originate from area and volume constraint forces arise from elastic properties of the membrane and cytoplasm of the cell. A lipid bilayer resists to expansion of its area, but only little to shear forces, which can be expressed by the force magnitude:

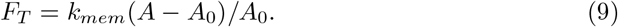

Here *k_mem_* is the area compression stiffness and *A*_0_, *A* are the reference and the current surface areas of the cell, respectively. These can be obtained by summing all the individual triangle areas, i.e. *A* = Σ_*k*_ *a_k_* of the cell surface, where *a_k_* is the surface area of a triangle *T_k_*, see Fig. 2B. Note that *A*_0_ is not necessarily constant as the cell can grow. The direction of the force **F**_*Tk*_ is from the barycenter of each triangle’s plane outwards [17]. The parameter *k_mem_* for area conservation forces in the membrane is related to the area bulk compression modulus *K_A_* by

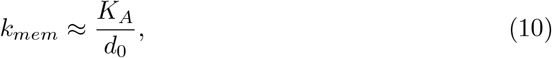

with *d*_0_ = ||**r**_0,*j*_ – **r**_0,*i*_|| the equilibrium length of triangle edge.

*Bending force - term (4) in Eq. 1:* The bending resistance from the cortex is incorporated by the rotational resistance of the hinges determined by two adjacent triangles *T*_1_ = {*ijk*} and *T*_2_ = {*ijl*}, see Fig. 2A. This permits definition of a bending moment *M*:

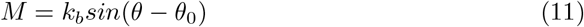

where *k_b_* is the bending constant, and *θ* is determined by the normals to the triangles **n**_*α*_, **n**_*β*_ by their scalar product (**n**_*α*_**n**_*β*_) = *cos*(*θ*). *θ*_0_ is the angle of spontaneous curvature. The moment *M* can be transformed to an equivalent force system **F**_*m,z*_ (*z* ∈ {*ijl*}) for the triangles *T*_1_ and *T*_2_ where here for *T*_1_ we can compute **F**_*m,i*_ = *M/l*_1_**n**_1_ using *l*_1_ as the distance between the hinge of the triangle pair and the point *i*, and similar expression for **F**_*m,l*_. The forces working on nodes *j, k* must fullfill **F**_*m,i*_ + **F**_*m,j*_ + **F**_*m,k*_ + **F**_*m,l*_ = 0 to conserve total momentum. The bending stiffness of the cortex is approximated by

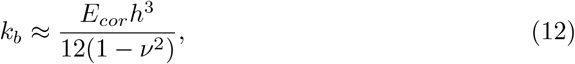

where *ν* is the Poisson’s ratio of the cortex.

*Volume force - term (5) in Eq. 1:* The volume change of the cell depends on the applied pressure and the cell bulk modulus *K_V_*. The compressibility of the cell depends on volume fraction of water in the cytosol, the CSK volume fraction and structure, and the compressibility of the organelles. In addition, it may be influenced by the permeability of the plasma membrane for water, the presence of caveolae [44], and active responses in the cell. We calculate the internal pressure in a cell by the logarithmic strain for volume change:

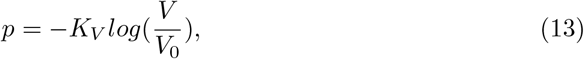

whereby *V* is the actual volume and *V*_0_ is the reference volume i.e., the volume of the cell not subject to compression forces. For small deviations of *V* from *V*_0_, *p ≈ K_V_* (*V − V*_0_)*/V*_0_. Within our model the volume *V* of the cell is computed summing up the volumes of the individual tetrahedra that build up the cell. The force *F_vol,i_* on node *i* is obtained by multiplying the pressure with the Voronoi area of node *i* (see [45]) and has a direction along the curvature vector **R**_*i*_ computed for that node (see contact model).

Eq. 13 expresses isotropic compression only. As *K_V_* ≫ *E_cor_ h_cor_ /R_c_* (see [46]), *K_V_* controls the overall compressibility of the cell while the mechanical stiffness of the cortex plays a minor role herein.

In case the internal CSK would be explicitly represented by internal structural elements not considered in the simulations of this work, those elements would contribute to both volume compression and shear forces.

*Cell-cell contact model forces - terms (8) in Eq. 1:* Whereas in a CBM, cells interact through central forces described by (modified) Hertz or JKR theory for adhesive spheres, in DCM the interaction forces need to be defined for each node individually. This can be achieved by pairwise potential functions (Van der Waals, Morse) between nodes that mimic the effect of short range repulsion and long ranged attraction forces of molecules [15, 16, 22, 47]. While straightforward to implement, this approach poses some problems with respect to the scalability and calibration of the parameters for these potentials. The approach followed in this work is different and adopts the method of Odenthal et al. (2013) [17, 48] which has been successfully applied to predict red blood cell spreading dynamics on surfaces. Here, we assign each triangle of the cell surface with a circumscribing sphere reflecting the local curvature. Two triangles belonging to different cells can interact by collision of their assigned spheres. To compute the magnitude of these interactions, we use Maugis-Dugdale theory. The Maugis-Dugdale theory for adhering bodies is a generalization of the JKR theory for spheres [49]. This theory captures the full range between the Derjaguin-Muller-Toporov (DMT) zone of long reaching adhesive forces of an soft homogeneous isotropic elastic sphere and small adhesive deformations of the Johnson-Kendall-Roberts (JKR) limit of a hard homogeneous isotropic elastic sphere of short interaction ranges.

In our model, Maugis-Dugdale theory is applied to a discrete system of triangles, which constitute the cell surface. This assumes a quasi-continuous distribution of cadherin bonds at the cell contact area^2^. As the cell curvature is not constant as it would be for a perfect sphere, we must locally estimate the curvature from the triangulated structure (see Fig. 2C) using the Laplace-Beltrami operator [17, 45, 48], which is a approximation function to associate a mean curvature vector to a discretized surface or boundary curve as alternative to approximation by the angle between the surface normals [51]. In this way every triangle of the cell is associated with a local curvature vector **R**_*i*_ for which a circumscribing sphere can be defined. An interaction between two cells defines several pairwise triangle-triangle interactions (*T, T′*). One pair of triangles define a pair of circumscribing spheres (*C*_1_, *C*_2_) and a common contact plane to which there triangles are projected (Fig. 2C, dashed red line).

The local stress which depends on the radius of both spheres and the adhesion energy, can then be computed using the adhesive Maugis-Dugdale stress component:

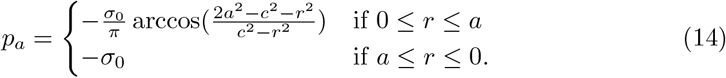

In this formula, *r* is the distance from the contact point, *a* is the effective contact radius (pure Hertz contact) and *c* is the radius of the adhesive zone (see Fig. 2, bottom). We then compute the Tabor coefficient [52]:

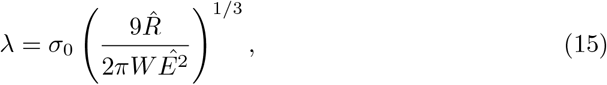

where 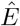 and 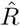 are the reduced elastic moduli and radii of the two objects in contact^3^. The tension *σ*_0_ is the maximum adhesive tension from a Lennard-Jones potential and is related to the adhesion energy *W* by *W* = *h*_0_*σ*_0_. Here *h*_0_ is the typical effective adhesive range that reflects the attractive cut-off distance between the bodies. We set *h*_0_ = 2 10^*−*8^ m in all the simulations [17, 53]. Following, *c* can be computed from *m* = *c/a* for which holds:

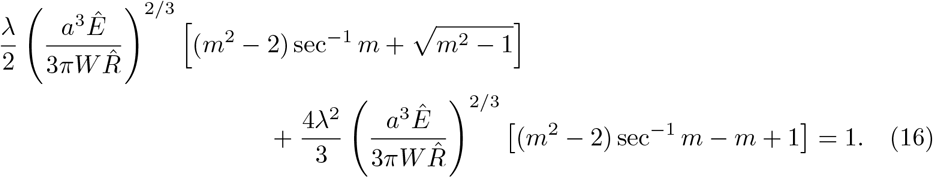

On the other hand, the repulsive part is given by the Hertz pressure:

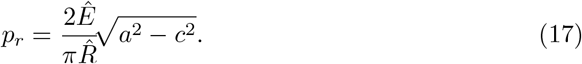

The total stress distribution between the two spheres is thus *p_a_* + *p_r_*. The total interaction force **F**_*adh,i*_ + **F**_*rep,i*_ between a pair of triangles is obtained by integrating this stress using standard Gauss quadrature rules, on the surface area common of the two triangles that have been projected on the contact plane. Once the force is known, it is distributed onto the nodes of both triangles, with the total force on the first triangle opposite in sign to the one of the second, to conserve total momentum. Note that for each triangle which has a contact area *A* with another triangle, any node associated to this triangle acquires a contact area *A/*3.

Importantly, this interaction model also permits to simulate interactions between a triangulated body and a smooth surface such as a sphere or plane having fixed curvature. In such case the contact is simply between the sphere assigned to a triangle, and the sphere that represents the object as a whole. This allows to implement an relatively simple algorithm that defines the handshake between a DCM and a CBM (see example Fig. 2D). For the full description of the model, see [17, 48].

*Cell migration force - term (9) in Eq. 1:* The migration force **F**^*mig*^ is usually an active force, representing the random micro-motility of a cell. Migration of cells involves complex mechanisms such as filopodia formation and cell contractility, and may be modeled as such (see e.g. [54, 55]), but here we do not go into such detail and lump these into one net force which is distributed to the nodes the cell. In absence of influences that impose a certain direction or persistence, it is commonly assumed that the migration force is stochastic, formally resulting in **F**^*mig*^ = **F**^*ran*^, with 〈**F**^*ran*^ 〉 = **0**, and 〈**F**^*ran*^(*t*) ⊗ **F**^*ran*^(*t*′)〉 = **M***δ*(*t – t′*), where **M** is an amplitude 3 × 3 matrix and relates to the diffusion tensor **D** of the cell. As cell migration is active, depending on the local matrix density and orientation of matrix fibers, the autocorrelation amplitude matrix **M** can a priori not be assumed to follow a fluctuation-dissipation (FD) theorem. However, “measuring” the position of a cell in the simulations the position autocorrelation function might be experimentally used to determine the diffusion tensor using 〈((**r**(*t* + *τ*) – **r**(*t*)) ⊗ (**r**(*t* + *τ*) – **r**(*t*))〉 = 6**D***τ*, and **M** be calibrated such that the numerical solution of the equation of motion for the cell position reproduces the experimental result for the position-autocorrelation function. For example, in a homogeneous environment **M** can be casted into a form formally equivalent to the FD-theorem, leading to an *k_B_T*-equivalent for cellular systems, that is controlled by the cell itself [13, 56].

On the other hand, if cells migrate in response to a morphogen gradient, an additional directed force **F**^*mor*^ into the direction of the concentration gradient of the morphogen may occur. The total migration force for the cell reads than **F**^*mig*^ = **F**^*ran*^ + **F**^*mor*^. In the liver lobule example simulations (Model III in section 3.3) we assume that only “leader” cells have the capability of directed migration. These are the cells in the lobule that are located directly at the border to the pericentral necrotic lesion.

Note, we assume here momentum transfer to the ECM by the ECM friction and active micro-motility term but we do not model the ECM explicitly.

#### 2.1.2 Cell growth and mitosis

During progression in the cycle, a cell grows by acquires dry mass and water, eventually doubling its volume. The inter-phase adding up the *G*_1_, *S* and *G*_2_ phase and mitosis phase may be described by an increase of the radius of the cell. We assume here that for a cell in free suspension during every the growth stage, the reference volume of the cell *V*_0_ gradually increases and is updated according to:

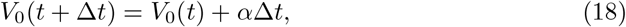

where *α* is the growth rate for simplicity assumed to be constant during the cycle, which can be justified if we look at timescales much larger that one cell division. *α* is chosen such that the volume of a cell doubles in the experimentally observed cell cycle duration. Cell growth and division in the CBM (see e.g. [6]) is straightforward to implement, but involved for the DCM requiring a multi-step procedure.

*Cell growth:* During cell growth, the volume of the cell is increasing (Fig. 3A-B). We model this by an update of the reference volume *V*_0_ following Eq. 13, update of the reference values of the cell surface area, and update of the spring constants. The reference triangle area *A*_0_ for each triangle, and spring length for each viscoelastic element are recomputed every timestep ∆*t* according to *A*_0_(*V*_0_(*t* + ∆*t*)*/V*_0_)^2*/*3^ and *l*_0_(*V*_0_(*t* + ∆*t*)*/V*_0_)^1*/*3^, respectively. The coordinated update of *V*_0_, *A*_0_ and *l*_0_ is necessary to ensure that no additional volume penalty forces or other artifacts are generated during growth. Note that at this momement, growth inhibition due to excessive external mechanical stress (e.g. [56]) can be easily included in our model, but in the scope of this paper we do not consider it further.

*Cell death:* When a cell dies, it can be either removed instantly from the simulation, or gradually shrink (lysis). Algorithm-wise, lysis can be regarded as the inverse of the growth process. However, during lysis the mechanical parameters of the cell may change. The process may continue until a certain cell volume has been reached, below which the cell cannot longer shrink. Finally, a lysed cell may be removed completely from the simulation (by e.g. phagocytosis).

*Cell division:* During cytokinesis, the continuous shrinking of the contractile ring, together with the separation of the mitotic spindle, gradually creates the new daughter cells. After mitosis the cell has split up in two adhering daughter. Such a process of cell division in 2D deformable cells has been previously described (e.g. [16]). In a first attempt we implemented such an algorithm in 3D but it turned out to be prone to numerical instabilities in 3D, as triangles on the side of the contractile ring tended to be extremely stretched while nodes accumulated at the position of the contractile ring. Hence, numerically stable simulation of these processes in DCM would require a complex re-meshing process for the surface in 3D, leaving the local stresses unchanged but avoiding numerical instabilities. An implementation of these steps turned out to be not only challenging, but also computing time consuming.

As we are merely interested in long term effects (i.e. several hundreds of cell divisions), and as the cytokinesis is a short process compared to the duration of the entire cell cycle, we avoid these particular tedious intermediate steps in our model, and directly create two new adhering cells that are within the shape of the mother cells just before its division as pictured in Fig. 3.

First, a division plane is chosen, which determines the direction into which the cells divide. The label of the division plane on the surface of the mother cell can be associated with the position of the contractile ring. The orientation of this plane may be chosen randomly or into a preferred direction, and splits the mother cells into two halves each bounded by part of the surface of the mother cell and the division plane (see Fig.3B-C); note that the mother cell in the figure is spherically shaped, but the algorithm works for arbitrary cell shapes). Then, the centers of the two future daughter cells on both sides of the plane are computed as the two centers of mass of the nodes that form each of the two halves, and each of those two centers of mass is associated with the center of a new spherically shaped daughter cell (Fig. 3C). The radii of the daughter cells are chosen such that they are both contained by the mother cell. To ensure the daughter cells are not overlapping at this stage, those nodes that would overlap the division plane are projected back on the division plane. Each of the daughter cells has now as border approximately a half of the mother cell envelop and the division plane that it shares with the other daughter cell. At this stage however, the radii (and volumes) of the daughter cells are not yet those they each should have, i.e. half of the volume of the mother cell.

To achieve this, a *sub-simulation*^4^ is performed where the volume and strut lengths of the viscoelastic network are reset to their reference values for a cell half the size of the mother cell. During the sub-simulation we artificially set all friction coefficients of the daughter cells to very small values. This will “inflate” the two daughter cells rapidly. On the other hand, we momentarily freeze the positions of the nodes of the mother cell. As a consequence, the two cells will rapidly adapt their shapes to the limiting shape of the mother cell “cocoon” and the division plane. The repelling interactions between the triangulated envelop of the mother cell and daughter cells ensure that the latter stay inside. We thus arrive at this step at a system of three triangulated cells: one fixed mother cell, and two encapsulated daughter cells with forming a shape approximately that of mother cell just before cell division.

After the sub-simulation, all the friction coefficients are reset to their normal values, the division plane is discarded and the daughter cells start adhering to each other because the mutual adhesion forces are invoked. The mother cell envelop is removed from the simulation and the daughter cells interact again with their environment. The system can relax slowly towards a mechanical equilibrium Fig. 3D over a time span equal to the mitosis phase duration.

### 2.2 Center Based Model (CBM)

Center-Based Models (CBM) are well established modeling approaches where cells have the same features as in DCM but the cell shape is approximated as a simple geometrical object. The precise cell shape is not explicitly modeled but only captured in a statistical sense. The cells are represented by homogeneous elastic and adhesive spheres (see Fig. 1A, bottom), and interact through pair-wise forces (e.g. Hertz, JKR), which are computed from a virtual overlap of both cell geometries. The equation of motion is similar to that of DCMs, yet the forces are here directly applied to the cell-centers (for details see section 2.3). During division, two new cells are created next to each other that replace the mother cell. In our work we will use the CBM for comparative runs in the simulations for liver lobule regeneration [4], see section 3.3.

### 2.3 Forces, equations of motion and cell division

We here only briefly recapitulate the basic components of CBM: for more detailed information, we refer to literature (see e.g. [5, 13, 56]). In CBM, cells are usually represented as spheres. The equation of motion for a cell *i* reads:

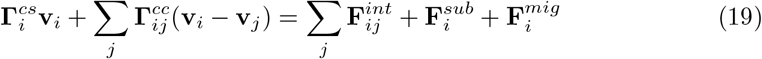

For the same reasons as for the DCM, inertia is neglected as cell movement occurs. The first term on the lhs. denotes friction forces of cell with the substrate or extracellular matrix, the 2nd term cell-cell or cell-sinusoid friction forces. Sinusoids are modeled in this work as immobile chain of slightly deformable spheres (hence **v**_*j*_ = 0 for sinusoidal elements *j*) with the radius equal to the sinusoidal radius found experimentally [57]. For cell-cell friction, the velocities **v**_*j*_ will generally be not zero. 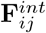 denotes interaction forces on cell *i* from repulsion or adhesion with other cells *j* or static blood vessel cells (see section 3.3). The force 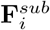 reflects adhesive/repulsive interactions with a flat substrate or wall. 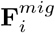 denotes the total migration force which has a random part and may also have a directed term (see explanation DCM). The friction terms involve tensors for the cell-cell friction 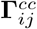 and cell-substrate friction 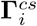. The friction tensors are computed in the same way as for the DCM, with the nodes of the DCM being replaced by the actual cell centers in the CBM [56].

The cells in CBM interact by pairwise potentials having a repulsive and adhesive part, which are characterized by a function of the geometrical overlap *δ_ij_*. As in [58] we assume here that cell adhesion forces can be described by Johnson-Kendal-Roberts (JKR) model, approximating cells by isotropic homogeneous sticky elastic bodies that are moderately deformed if pressed against each other. The interaction force is computed by

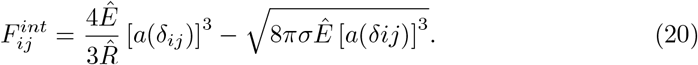

The contact radius *a* in Eq. 20 allows to compute the cell-cell contact area, and can be obtained (from [59]):

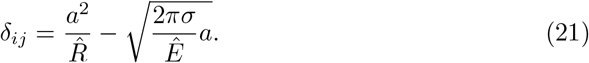

In the latter equations, 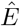 and 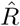 are defined as

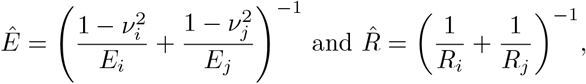

with *E_i_* and *E_j_* being the Young’s moduli, *ν_i_* and *ν_j_* the Poisson numbers and *R_i_* and *R_j_* the radii of the cells *i* and *j*, respectively. Note that the Young moduli in the center-based model should be chosen such that it corresponds as much as possible to the elastic properties of the DCM. In particular, we warrant here that for a CBM’s Young modulus and Poisson’s ratio, *E* = *K_V_ /*3(1 − 2*ν*) where *K_V_* is the compression modulus of the DCM cell (section 2.1). Note that for a CBM, the elastic properties of the cortex and membrane a priori cannot be identified.

During the cycle the intrinsic volume of a mother cell doubles before it splits into two daughter cells. The intrinsic volume is defined by the volume of the cell if it would be undeformed and uncompressed. Its true volume in case it interacts with other cells or structures cannot be exactly tracked as the CBM does mostly not permit to calculate the volume of cells interacting with other cells. Like in the DCM we are assuming constant growth rate during the cell cycle and the intrinsic volume *V_i_* of cell *i* is updated every timestep ∆*t* as in Eq. 18.

If the cell passed a critical volume *V_crit_*, the cell undergoes mitosis and two new cells with volume *V*_0,*i*_/2 are created. A simple of the division algorithm consists of placing directly two smaller daughter cells in the space originally filled by the mother cell at the end of the interphase [12, 60]. When the two daughter cells are created, the JKR interaction force will push away the two daughter cells until mechanical equilibrium is reached. If the space filled by the mother cell is small, which is often the case for cells in the interior of a cell population, the local interaction forces occurring after replacing the mother cell by two spherical daughters, can adopt large (unphysiological) values leading to unrealistic large cell displacements. This might be circumvented by intermediately reducing the forces between the daughter cells (see below). Alternatively, cells could in small steps be deformed during cytokinesis into dumbbells before splitting [4]. In this work we pursue the simpler approach as it resembles the cell division algorithm we use for the DCM.

#### 2.3.1 Corrections to JKR contact forces

As mentioned above, center-based models suffer from a number of major artifacts, often ignored in simulations. The most striking is that common pair-wise contact forces (type Hertz, JKR,..) and contact areas become largely inaccurate when cells are densely packed (jammed) and compressed [13]. The origin of this problem is that these forces are defined pairwise and exclude the contributions from other interactions. As a consequence, even an incompressible cell characterized by Poisson ratio *ν* = 0.5 in the JKR force model (Eq. 20) may be compressed if surrounding cells are pushed towards it. In such a situation, cell volume and cell-cell contact area are only poorly approximated by the JKR force model.

Voronoi tessalation of the positions of the cells can estimate the individual cell volumes (and hence predict realistic pressure forces) [12, 61]. However, in the attempt to correct the unrealistic contact forces in the center-based model upon large compression forces, we here propose a simple calibration step in which we use DCM simulations to estimate contact forces between cells. The DCM does not have the aforementioned problem because the shape and cell volume are exactly determined. As such, there is no notion of geometrical overlap in DCMs. A small clump of DCM cells is compressed quasi-statically while monitoring the contact forces as function of the distances between the cell centers (see section 6.1, Supplementary Material Fig. 12B, and Video 5 in the Supplementary Material). During compression, a stiffening of the contacts can be clearly observed. Performing an equivalent experiment with CBM using the JKR law results in a significant underestimation of the contact forces (Fig. 12B). To correct this, we keep the original JKR contact law form but modify the apparent modulus 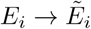 of the cell as the cells approach each other, by gradually increasing it in Eq. 20 as the cells get more packed. The challenge lies in determining 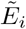 as a function of the compactness of the spheroid. We try to estimate the degree of packing around one cell by using the distances between that cell and its neighbors, introducing a function that depends on the *local* average of the distances *d* = Σ*_j_ d_ij_ /N* (N is the number of contacts) to each of the contacting cells *j*, noticing that the cell-cell distances differ only very slightly. The simulated force curves of the DCM could well be reproduced with the CBM for the following simple polynomial function of 4th order:

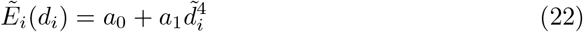

in where *a*_0_ = *E_i_* and, *a*_1_ is a fitting constant. The best fit to the DCM contact force is shown in Fig. 12A, with *a*_1_ = 4 10^6^. However, we note that the contact stiffness generally depends on the number of neighbors a cell has. In this experiment, every cell had initially 8 contacting neighbors, yet this number evolves to approximately 12 as the compression progresses, which is what would be expected from the Euler theorem for volume tesselation.

A side effect of this calibration method is that high repulsive forces may arise during cell division, when two new cells are created and positioned close to each other. To limit these effects, we apply the above formula in a such way, such that the contact stiffness becomes gradually larger over a time period just after division, which we choose about ∼ 1*/*10 of the cell cycle (roughly the duration of mitosis phase; however, smaller durations would also be possible), reaching the maximal value after this period. This is similar as what has been used in Galle et al. [60] to avoid repulsion force cues, and ensures that cells will not separate abruptly during division. The period gives time to the cells to evolve to a local mechanical equilibrium, and can be seen as the analogue of the relaxation period in the division algorithm in the deformable cell model (see section 2.1.2). We have verified that small variations on this relaxation period did not influence the simulations results significantly.

## 3 Results

### 3.1 Single cell experiments

#### 3.1.1 Calibration of DCM parameters from optical stretcher experiments

Classical experiments such as optical stretchers, optical tweezers, and micro-pipetting techniques are used to observe the mechanical behavior of individual cells in ground state and can be used to estimate the physical range of the DCM viscoelastic network model parameters (see e.g. [17, 26, 32]). Here, we choose the optical stretcher experiment, which is mimicked *in silico* with a DCM. In the optical stretcher, two laser beams in opposite direction and faced to each other trap a cell in suspension (see Fig. 4A). The diffracted laser beams exert a surface force on the cell’s membrane and cortex, deforming it towards the beam direction. Increasing the laser power yields a higher optical stretching force. At the same time the deformation along two perpendicular directions is measured using image analysis, yielding information on cell shape. We refer to refs. [36, 62, 63] for more details of the setup and conditions in such an experiment.

**Fig 4.**
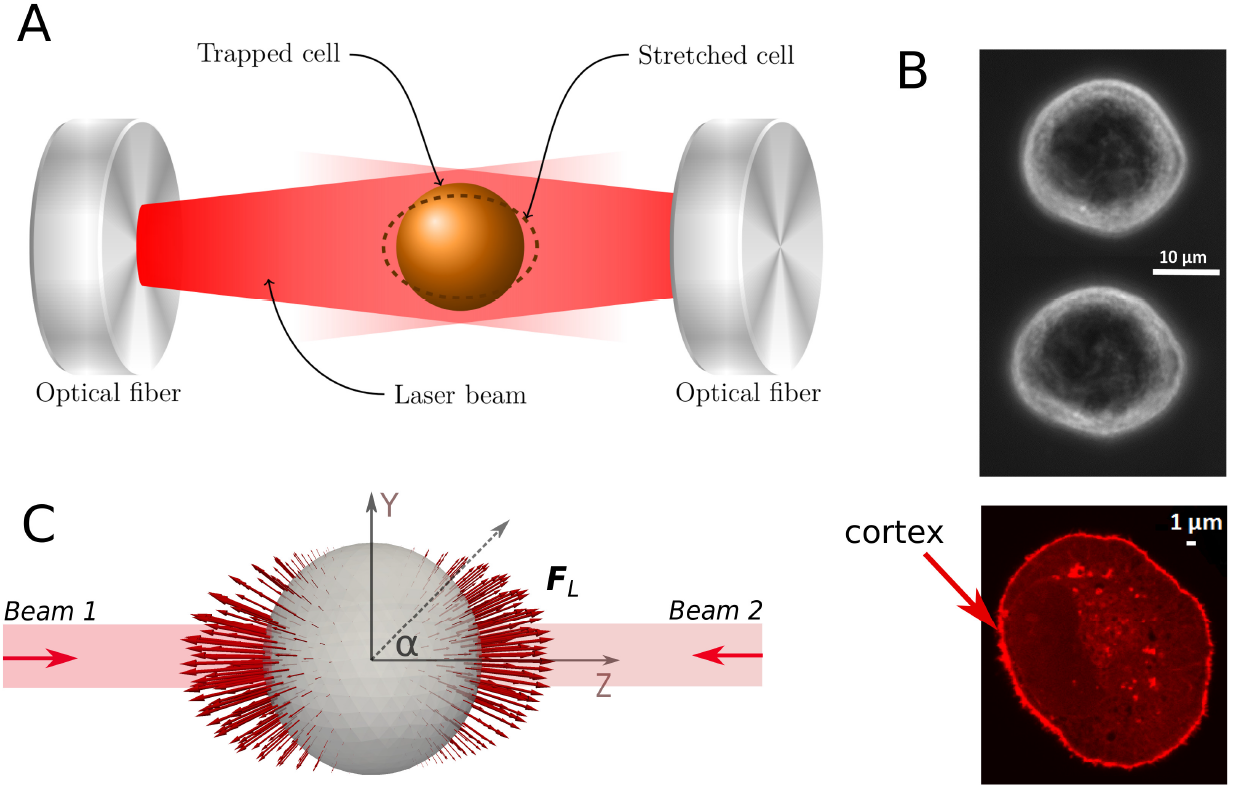
(A) Sketch of the experimental setup with a trapped cell in an optical stretcher. (B) Top: image of unstreched and stretched MDA-MB-231 cell. Bottom: actin stained image of a MDA-MB-231 cell indicating cell cortex actin cytoskeleton. (C) Surface forces profile due to laser is visualized (red arrows) on a triangulated surface of the deformable cell. The Z-axis (long axis) is aligned with the laser beam, whereas the X-and the Y-axis give the direction of the short axes during deformation. **F**_*L*_ is the nodal force induced by the laser power.

New data of optical stretcher experiment was generated using MDA-MB-231 breast cancer cells. These cells had an average radius of 8.8 *±* 1.3 *µ*m (N= 100 samples). Actin staining further revealed average approximate cortex thickness of *h_cor_* ∼ 500 *µ*m (Fig. 4B). Hepatocytes were not amenable to optical stretcher experiments.

In each experiment the laser beams were applied during a time interval of a few seconds, in which the cell continues to deform. Thereafter, the cell relaxation behavior period was monitored for several seconds. The long axis (Z) and short axis (Y) lengths changes over time are derived from analyzing the images (see data Fig. 5; the X-axis and Y-axis are by rotation symmetry with regard to the Z-axis assumed to be equally long). Two laser powers were considered (*P*_0_ = 900 mW and *P*_0_ = 1100 mW), with an applied stretching time of two seconds and a monitoring of the relaxation behavior of two seconds. Because of the large biological variability, for each individual experiment the measurement has been performed with a minimum of about 100 cells.

**Fig 5.**
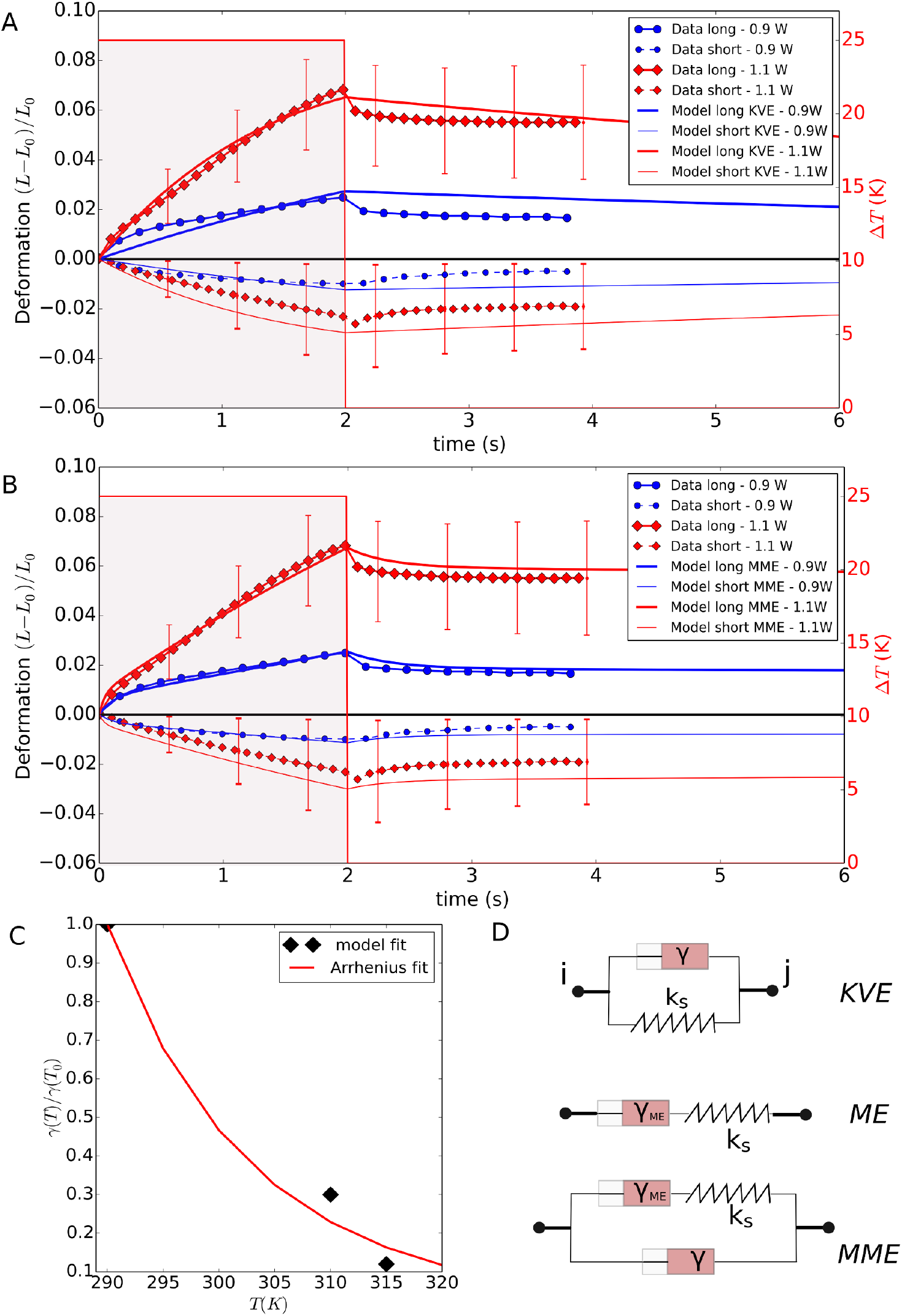
(A-B) Comparison of the simulated MDA-MB-231 cell deformation with the experimental data in an optical stretcher, using laser powers of 900 mW and 1100 mW. (A) The model was first fitted using a Kelvin-Voigt model (KVE) and assuming temperature dependent friction coefficients for the cytoskeleton. (B) The model then was fitted assuming a modified Maxwell model (MME) but using the same temperature dependence as for KVE. For sake of clarity, the error bars are only shown for some data points, and only for the *P*_0_ = 1100 mW case. The errors bars on the data are quite large, indicating a high biological variability. (C) Temperature dependence of damping coefficients, using model fit for DCM, compared to Arrhenius’ law using an activation energy close to the value reported in ref. [64]. (D) Schematic representation of nodal force elements with springs and dampers. KVE: Kelvin-Voigt element, ME: Maxwell element, MME: modified Maxwell element.

In a first step we identified some parameter and/or their ranges by comparison to published references.

An initial spherical deformable cell was created with a total of *N* = 642 nodes (test runs indicated that further spatial refinement was not required). The cells were immersed in a medium with viscosity *µ* being close to that one of water. We approximate the cell-medium friction coefficients for each note needed to determine term (3) in Eq. (1) by *γ_ns_* = *γ_liq_ /N* = 6*πµR/N*. This approximation ensures that for a spherical cell of radius *R_c_* modeled by *N* nodes at its surface, the friction coefficient is precisely as predicted by the Stokes equation.

Determination of the force terms (4) and (5) in Eq. 1 require determination the elastic modulus of the cell cortex. Because we did not have information on the elastic properties of the MDA-MB-231 cells, we adopted a nominal value for the elastic modulus of the cell cortex *E_cor_* ∼ 1 kPa based on different cell types (see [46, 65]). Applying Eq. 7 and Eq. 12, which relate the elastic moduli to the parameters of the coarse grained model of the CSK, the nominal cortical stiffness constants in the model are thus *k_s_* ∼ 4 10^*−*4^ N*/*m, *k_b_* ∼ 1.10^*−*17^ Nm (see section 2.1, Eq. 3 and Eq. 12). To determine the compression force of the cell represented by term (6) in Eq. (1), we need to know the cell compression modulus *K_V_*. The cell compressibility is still subject for debate. For instance, Delarue et al. [66, 67] conclude from experiments with growing spheroids under pressure that cells are compressible with compression moduli of the order of 10 *kPa*. On the other hand, the Monnier et al. [42] find individual cell compression moduli of several orders of magnitude higher (1 MPa) than the one reported above. Yet, Monnier et al. have measured this over short time period and accordingly they further state in their paper that on longer timescales, the compression modulus might differ from that value may due to adaptation of the cell. In the work Tinevez et al. [46] the cytoplasm compression modulus is estimated as +/- 2500*Pa*. Despite this is not the compression modulus of the whole cell, it indicates that if cells are able to expel water through the aquaporins on longer timescales, this may be a good estimate of *K_V_*. We here adopted this value but we note that in the simulations for the optical stretcher, we did not find any significant influence on the results during the time course of the experiment when *K_V_* was varied within 1000 Pa to 100000 Pa.

Term (7) of Eq. (1) represents the effect of bilayer compression modulus *K_A_* (see Eq. 9). For pure lipid bilayers about 0.2 N*/*m have been reported, while in case of a plasma membrane of a cell, much lower values have been measured, likely due to invaginations and inclusions of other molecules in a regular cell membrane. In our model, we chose *K_A_* = 0.8 mN*/*m [65].

In the work of Ananthakrishnan et al. [68] it was suggested that the cortical actin cytoskeleton is the main component determining the mechanical behavior of the cell. We follow this assumption, yet in order to simulate the experiment, we need know the applied surface force vectors on each node generated by the deflecting laser beams (see Fig. 4C). The calculation of this force profile is described in Appendix section 6.2.

In a second step we identify the type of the viscoelastic elements connecting the surface nodes of the model cells (Fig. 5D) by comparing the results of the model simulations with those of the optical stretcher experiments with MDA-MB-231 cells. The characteristic relaxation behavior excludes purely elastic springs (Fig. 5). we first consider Kelvin-Voigt elements (KVE) between the surface nodes (Fig. 5D). Kiessling et al. (2013) [64] reported thermoelastic phenomena, whereby an increased laser power causes a transient temperature rise, which in turn modifies the viscoelastic properties of the cell. A thermal analysis of the used setup similar as in [64] indicated an ambient temperature increase from 20 K to 25 K for a laser power from 900 mW to 1100 mW. The temperature rise and fall is very quick (∼ *ms*) in comparison to the duration of the experiment, hence latency effects can be neglected. Because of the thermoelastic effects, we assumed in the model that the viscous properties (i) are changed by the laser power during the stretching stage to (i.e. *γ* = *γ*(*T*)), and (ii) are all restored to their original value during release. A good fit for the *P* = 1100 mW was achieved by the values *k_s_* = 1.0 *·* 10^*−*4^ N*/*m, *k_b_* = 2 *·* 10^*−*17^ Nm as cortex elastic constants, *γ_||_* = *γ*_⊥_ = *γ*(*T*_0_) = 2.3 *·* 10^*−*4^ Ns*/*m as nominal friction coefficient (see Fig. 5A) and by a temperature dependence of the friction coefficients of *γ*(20)*/γ*(*T*_0_) = 0.29 and *γ*(25)*/γ*(*T*_0_) = 0.11. The fitted elastic constants for the springs and bending thus correspond well to those obtained from the nominal material properties (see above). We then tested the assumption that the viscosity would scale with temperature as in the Arrhenius law [64]:

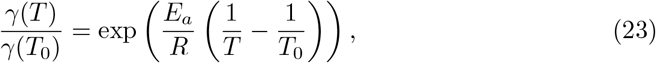

where *R* = 8.314 Jmol^*−*1^K^*−*1^ and *T_ref_* = 290 K. Using this equation we got an optimal fit with the model calibrated friction coefficients (see Fig. 5C) using the value *E_a_* = 55 KJmol^*−*1^K^*−*1^, which is not too far to *E_a_* = 74 KJmol^*−*1^K^*−*1^ derived in [64]. Neglecting thermoelastic effects, no simultaneous fit of the experimental results for both laser powers could be obtained: if the KVE model is calibrated such that an optimal fit is obtained for *P* = 1100 mW, then using the same constants we get a significant overestimation of the deformation for *P* = 900 mW, and vice versa (result not shown).

Although the stretching and the relaxation behavior are both qualitatively well captured if thermoelastic effects are accounted for, there is (i) a restoring effect of the cell shape in the simulations (solid like behavior) that is not observed in the time course of the experiment, and (ii) the short axis deformation slightly exceeds the experimental one. This latter deviation (point (i)) might be a consequence of missing out an explicit representation of the internal cytoskeleton in the model (i.e. see [69]), which might here result in a higher resistance to a movement of the cell surface perpendicular to the laser axis, but including an explicit representation of the cytoskeleton is not expected to remove the significant deviations of the model during the relaxation stage (point (i)). For this reason, we capture a possible contribution from the fluid like behavior of the cortical cytoskeleton, see e.g. [65], by adding an additional friction element in series with the spring in the Kelvin-Voigt element (with the choice of *γ*(*T*_0_) = 10^*−*5^ Ns*/*m), resulting in a modified Maxwell model^5^ (MME), see Fig. 5D. An optimal fit for the *P* = 1100 mW is now achieved using the elastic constants *k_s_* = 5 10^*−*4^ N*/*m, *k_b_* = 2 10^*−*17^ Nm, but with *γ*(*T*_0_) = 3 10^*−*5^ Ns*/*m and *γ_M E_* (*T*_0_) = 3 10^*−*3^ Ns*/*m as overall damping parameter and for the Maxwell damping respectively (Fig. 5B). The temperature dependence for the friction coefficients was chosen as obtained in the KVE model (Eq. 23). The two fits capture the relaxation behavior overall well and much better than the KVE model, with an apparent plastic deformation keeping the cell shape from its original value.

In conclusion, in this section we have identified the basic biomechanical material properties of the DCM, namely the viscoelastic elements and its parameters, to quantitatively mimic optical stretcher experimental results. Our simulations indicate that the laser induced stretching and the subsequent fluid like relaxation behavior after the stretching can be best mimicked assuming a modified Maxwell model for the elements (MMEs) representing the cortical cytoskeleton, with the additional assumption that the viscous properties decrease with increasing temperature as a result of the lasers. The KVE model overall results in less optimal fits but may still be useful in simulations, because the behavior of the cells, on longer timescales (minutes to hours) after the experiment is not a priori known and may exhibit active responses, possibly leading to a restoring of the original cell shape [70].

#### 3.1.2 Verification of cell adhesion forces in pull-off experiment

The DCM adhesion model used in this work has been able predict the correct red blood cell spreading area on a surface given a certain adhesion strength [17]. We have further validated this adhesion model on the level of the adhesive forces between two cells. In ref. [58] it was shown that JKR theory can be applied to living cells using micro pipette aspiration, providing a technique to determine the adhesion energy between cells experimentally. To verify that the resulting adhesive forces in our discretization are in the correct physical ranges, we have run several test simulations in which two adhered cells (with radius *R*_1_ = *R*_2_) are mechanically separated by applying opposite forces to them. The separating force (also called pull-off force) is a well-known quantity in JKR theory for soft adhesive spheres and reads

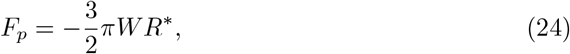

where *W* is the adhesion energy per surface area (sometimes called specific adhesion energy) and 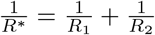. On the other hand, adhesion between stiff spheres is best described by Derjaguin-Muller-Toporov (DMT) theory [53], leading to a pull off force of *F_p_* = *−*2*πW R^∗^*. For vesicles, a theoretical analysis leads to *F_p_* = *−πW R* for *R*_1_ = *R*_2_ [53].

To measure the cell-cell adhesion force in our simulations, two cells in contact are first brought into contact and relaxed until they are in an equilibrium state. The adhesion energy is a genuine parameter in our model (see section 2.1). Then, a force equal in magnitude is applied on each node of both cells into the direction perpendicular to the cell-cell contact, whereby the force on one cell is exerted in opposite direction than the force on the other cell. The orientation is chosen such that the contact is under tensile stress (negative load). The cells then move away from each other until a new equilibrium is reached (Fig. 6). The force at which the cells just separate defines the pull-off force. By performing several simulations following a binary search algorithm, we estimate the pull-off force. The operation is done for increasing adhesive forces and depicted in Fig. 6C. This shows that the simulated pull-off forces are indeed very close to the theoretical values as calculated from JKR theory, DMT theory, and vesicle theory. Overall, we can assume that the implementation of the adhesion energy and the choice of parameters in the model reproduce realistic magnitudes for the contact force.

**Fig 6.**
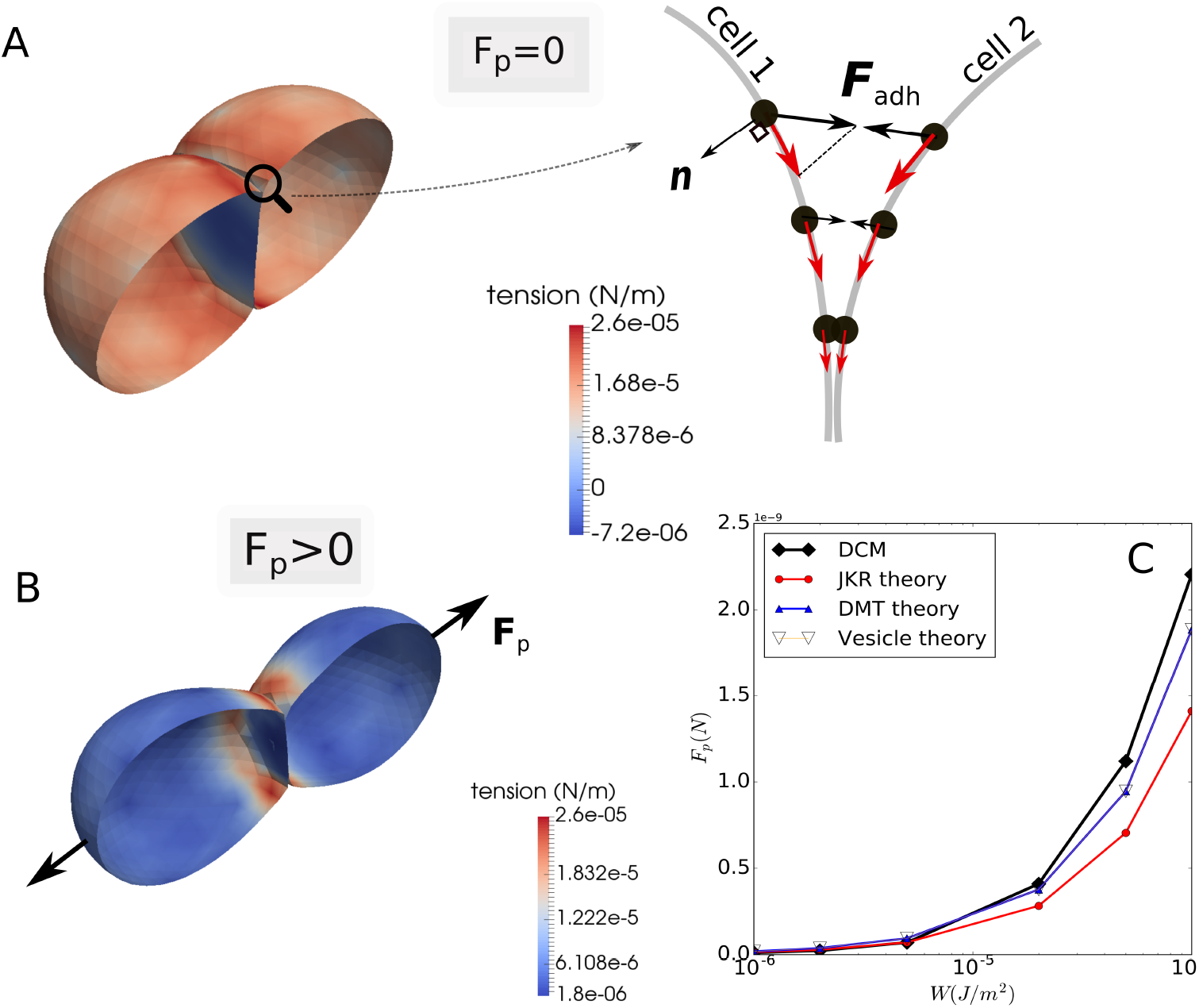
(A-B) Simulation snapshots of the experiment for determining pull-off force for adhering cells. The color coding is according to membrane stress. Stresses can be negative (compressive) in the common adhesive plane, indicated by the blue coloring (A, left). The 2D cartoon (A, right) shows the forces on the nodes **F**_*adh*_. The resulting nodal force (red arrows) along the cell boundaries point towards the contact center. (B) Cell deformation during the pulling with net force *F_p_*. (C) Absolute values of pull-off forces obtained as function of the adhesion energy per unit surface area: DCM simulations compared to JKR theory, DMT theory, and vesicle theory. Video 1 of this in silico experiment is provided in the SI 6.3.

A closer analysis of Fig. 6 (top) further reveals that tensions in the membrane (see Eq. 8) of the adhered cells in equilibrium are largely tensile except in the adhesive plane where they can be compressive. This is a direct consequence of the adhesive node-node forces at the boundary of a cell-cell contact, which generate forces (see red arrows in sketch, Fig. 6) on the nodes along the cell-cell interface towards the contact zone. This phenomenon is in agreement with experimental observations [71]. A very strong adhesion energy might even result in a buckling of the cell surfaces at the contact zone as we observed in test simulations, and as was suggested in [72].

### 3.2 Multicellular simulations: growth of small tumors and monolayers

In the past, multicellular spheroids (MCSs) and monolayers have often been used and tested as in-vitro models for tissues (see e.g. [73, 74]). Several authors have investigated the growth dynamics of these systems using agent-based models. In a number of communications, center-based models (CBM, see section 2.3) have given basic insight how cell growth is affected by hypoxia and contact inhibition [5, 12, 56]. As mentioned before, CBMs generally suffer restrictions, such as poor shape representation.

Here, we employ our deformable cell model including cell division with regard to the aforementioned systems. In the growth dynamics of multicellular assemblies, important parameters are the strength of cell-cell adhesion responsible for the formation of multiple cell-cell contacts, and viscous cell-cell friction. To determine the cell-cell friction coefficients, we have simulated the relaxation behavior in an experiment whereby a spheroid is first compressed and subsequently the compression released, see Appendix 6.1. For long compression times, typical relaxation times of ∼ 5 hours have been reported for such experiments [75, 76]. Hence the spheroid relaxation times are much longer than the short relaxation timescales of the cells in the stretcher experiment, indicating that cellular re-organization processes not relevant in the optical stretcher experiments play a major role on the time scale of multicellular spheroid relaxation. E.g., trypsinated cells in suspension round off indicating that on time scales much longer than those probed in optical stretcher experiments, the cell relaxation behavior is governed by processes restoring a spherical cell shape, which is not correctly captured by the modified Maxwell element mimicking short term relaxation (Fig. 5D). Instead of further complexifying the viscoelastic elements to capture both the short and long-term relaxation behavior at the expense of elevating the computing time requirement^6^, we here use the simple KV viscoelastic elements for the simulations at the time scale of cell growth and division. This allows to fit the internal friction coefficients and cell-cell friction coefficients such that the spheroid relaxation is approximately 5 hours without prolonging the computing time (see Fig. 12B).

Both MCSs and monolayer simulations start from a single cell growing and dividing unlimited with a cell cycle time of 24*h*. Each cell is represented by 162 nodes. For an overview of all the reference parameters, see Table 1. The simulations are run over several days, in which in total about 1000 cells are created (see time series of simulation snapshots in Fig. 7A, B for MCSs and monolayers, respectively). Two cases were simulated, one with the original specific adhesion energy, *W* = 10^*−*5^ J*/*m^2^, one with an increased specific adhesion energy, *W* = 8.10^*−*5^ J*/*m^2^. However the cell population sizes were too small to objectify differences in the tumor spheroid shape despite the spheroid with *W* = 8.10^*−*5^ J*/*m^2^) looks slightly more compact compared to the other case, where intermittently, small “branches” form (Fig. 7A, right). At this cell population size, the number of cells grows exponential in time in both cases (Fig. 7C).

**Table 1.**
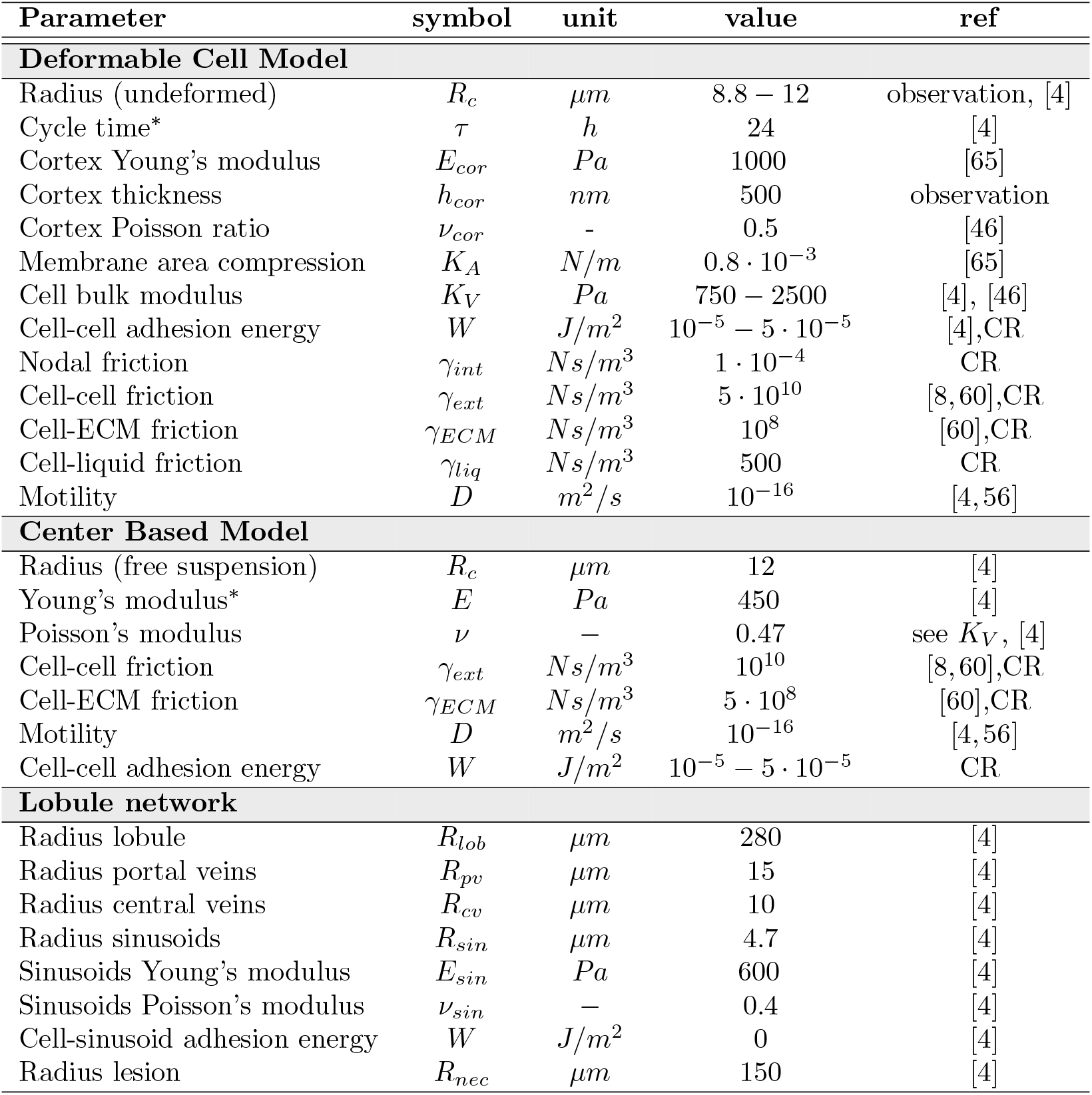
Nominal physical parameter values for the model. An (*) denotes parameter variability meaning that the individual cell parameters are picked from a Gaussian distribution with *±*10% on their mean value. CR: Calibration Runs.

**Fig 7.**
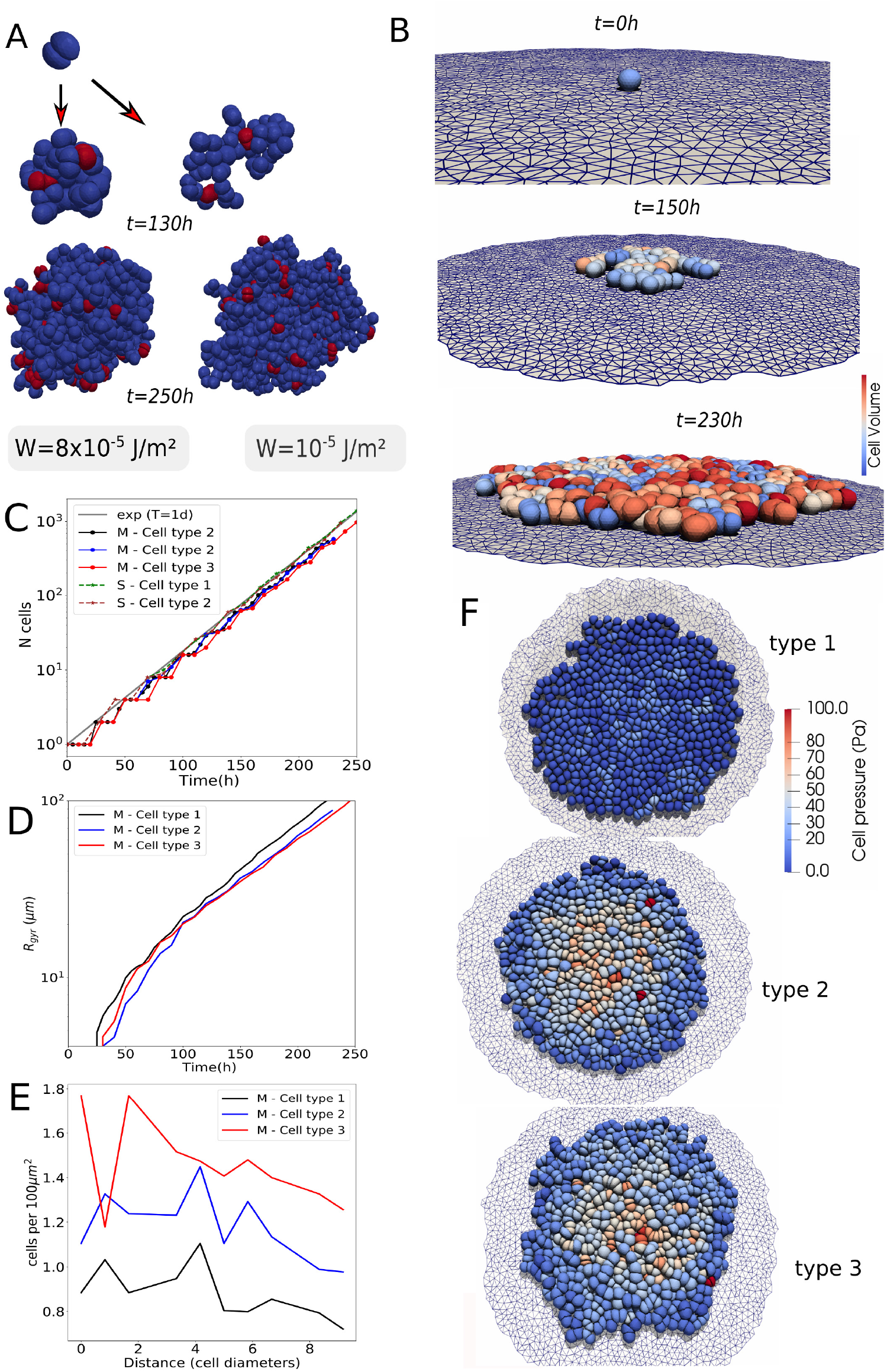
Simulation snapshots and time series for multicellular spheroid (MCS) and monolayer growth. (A) Spheroid growth with specific cell-cell adhesion energies *W* = 1.10^*−*5^ J*/*m^2^) and *W* = 8.10^*−*5^ J*/*m^2^). (B) Typical monolayer growth scenario (screenshots). (C-E) Kinetics of the cell population size *N* (*t*) (monolayers, MCS), radius of gyration *R_gyr_*(*t*) (monolayers) and area density profile (number of cells per unit of substrate area) *ρ*(*r, N* = 1000) (monolayers). (F) Visualization of spatial pressure distribution in multicellular pattern for type 1-3 monolayers (the color encodes internal cell pressure (red: high, blue: low pressure)). Parameters for monolayers: type 1: cell-cell specific adhesion energy *W* ≈ 0*J/m*^2^, cell-substrate adhesion energy *W* = 10^*−*5^*J/m*^2^, *D* = 10^*−*15^*m*^2^*/s*; type 2: cell-cell specific adhesion energy *W* = 10^*−*5^*J/m*^2^, cell-substrate adhesion en2e5rgy *W* = 10^*−*5^*J/m*^2^, *D* = 10^*−*16^*m*^2^*/s*; type 3: cell-cell specific adhesion energy *W* = 5 × 10^*−*5^*J/m*^2^, cell-substrate adhesion energy *W* = 10^*−*5^*J/m*^2^, *D* = 10^*−*16^*m*^2^*/s*. Video 2 of this in silico experiment is provided in the SI 6.3.

In a further step, we test our model for three different cases of monolayer growth. We first constructed a flat fixed adhesive surface, triangulated with a slightly larger mesh size than the one used for the cell surface triangulation. The simulation starts from one cell adhering to the surface (the reference adhesion energy per area is *W* = 10^*−*5^ J*/*m^2^). To study the effect of adhesive strength, we simulated monolayer growth scenarios for three different parameter combinations, denoted as type 1-3, with type 1: specific cell-cell adhesion energy *W* = 10^*−*16^*J/m*^2^, and relatively large diffusion constant *D* = 10^*−*15^*m*^2^*/s*; type 2: specific cell-cell adhesion energy *W* = 10^*−*5^*J/m*^2^, and diffusion constant *D* = 10^*−*16^*m*^2^*/s*; type 3: specific cell-cell adhesion energy *W* = 5 10^*−*5^*J/m*^2^, and diffusion constant *D* = 10^*−*16^*m*^2^*/s* (Fig. 7C-F). Visually no large difference between the spatial pattern for different parameter combinations can be seen. This is objectified by plotting the radius of gyration *R_gyr_*(*t*) for the three different parameter combinations, which is a measure for the spatial cell spread. It is defined as *R_gyr_* = 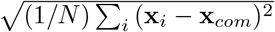, where **x**_*i*_ and **x**_*com*_ = (1*/N*) _*i*_ **x**_*i*_ are the center-of-mass positions of the individual cells and the whole multicellular cluster, respectively. There is no significant difference in *R_gyr_* for the three different types. All three populations grow exponentially fast during the simulation time period. However, the pressure profile differs: at a population size of 1000 cells, the pressure in the center of the monolayer is largest for type 3, the smallest for type 1 and intermediate for type 2. For type 1, the cell-cell adhesion energy is almost zero while the micro-motility is the largest, which results in a maximum relaxation of compressive stress and lower cell density (Fig. 7E). The specific cell-cell adhesion energy for type 3 cells is higher than for type 2 cells while all other parameters are the same, resulting in a higher adhesion and hence a higher compression (highly jammed state) in the interior of type 3 cell populations compared to the interior of type 2 populations (Fig. 7E). The spatial profile shows a decrease from the center to the border, in qualitative agreement with earlier observations in center-based model simulations (e.g. [60, 77]) but different from the center-based model. Contrary to the former studies, the cell volume and pressure in DCM simulations can be calculated more precisely.

In summary, these simulations show the potential of the model to analyze in detail forces, shape and pattern formation in monolayers and spheroids. We must emphasize that the aforementioned results are indicative and that much more and much longer simulations maybe required to come to strong conclusions.

### 3.3 Multicellular simulations: regeneration in a liver lobule

Finally we come back to the introductory example of the regenerating liver lobule after CCl4-induced peri-central damage, and study whether the regeneration process with the DCM model would lead to different results than obtained with the CBM model [4] The simulations are performed in a in statistically representative liver lobules, obtained by sampling of parameters quantifying the lobule architecture in confocal laser scanning micrographs (Hoehme et al., 2010).

A statistically representative lobule has an approximately hexagonal structure (Fig. 8A-B, XY plane) with a vertically oriented central vein in its center (oriented into the direction of the Z axis). The distance from the central vein to the lobule border (radius of the lobule) is about 9 cell diameters (Hammad et. al., 2014) [57]. A typical 3D volume data sample obtained by processing of optical sections from confocal micrographs has a height of three cell layers. Each lobule has about three portal triads localized in differently of the on average six corners of the lobule. A triad consists of a portal vein, carrying blood from the intestine to the lobule, a hepatic artery, transporting oxygenated blood from the aorta to the lobule, and a bilary duct, transporting bile away from the lobule towards the gall bladder. The blood flows through a network of sinusoids, fenestrated capillaries surrounded by hepatocytes and drains into the central vein, from where it is transported away from the liver. Between sinusoids and hepatocytes there is an about 0.5*µm* small space, named “space of Disse” filled with ECM that mechanically stabilizes the sinusoidal network. The liver lobule micro-architecture ensures a maximum exchange area between hepatocytes and sinusoids for metabolites, thereby promoting metabolisation in the hepatocytes. After overdose of the drugs CCl4 or acetaminophen (paracetamol, APAP) those cells expressing the Cytochrome P450 enzymes CYP2E1 and CYP1A2 metabolize these two drugs to NAPQI, which downstream causes cell death. In healthy liver, CYP-expressing enzymes are localized in hepatocytes within an area fraction of about 40% around the central vein. At sufficiently high doses of CCl4 (or acetaminophen) the hepatocytes are killed leaving a central lesion with debris with maximal size at about 1-2 days after administration of the drug. About two days after the generation of the lesion, hepatocytes start to divide, reaching closure of the necrotic lesion and regeneration of the hepatocyte mass after about 6 days [4]. The hepatocytes localized at and close to the border of the lesion proliferate by far the most, while proliferation is lower in the layers more distant to the lesion. Simulations with a center-based model revealed that cell proliferation alone, without directed migration of hepatocytes towards the central vein, is insufficient to explain the experimental observations [4]. The mechanical pressure exerted by proliferating cells on their neighbors leading to pushing of cells and squeezing them into the spaces between the sinusoidal network was insufficient to close the lesion within the experimentally observed time period. In center-based models (CBM), cells are by construction usually relatively rigid as only moderate deviations of cell shape from their equilibrium shape in isolation can be captured by that model type. Furthermore, forces in standard center-based models are defined pairwise lumping compression and deformation forces usually together (see also section 2.3.1). We here study if this rigidity might be responsible for the failure of regeneration in the experimentally observed time period.

**Fig 8.**
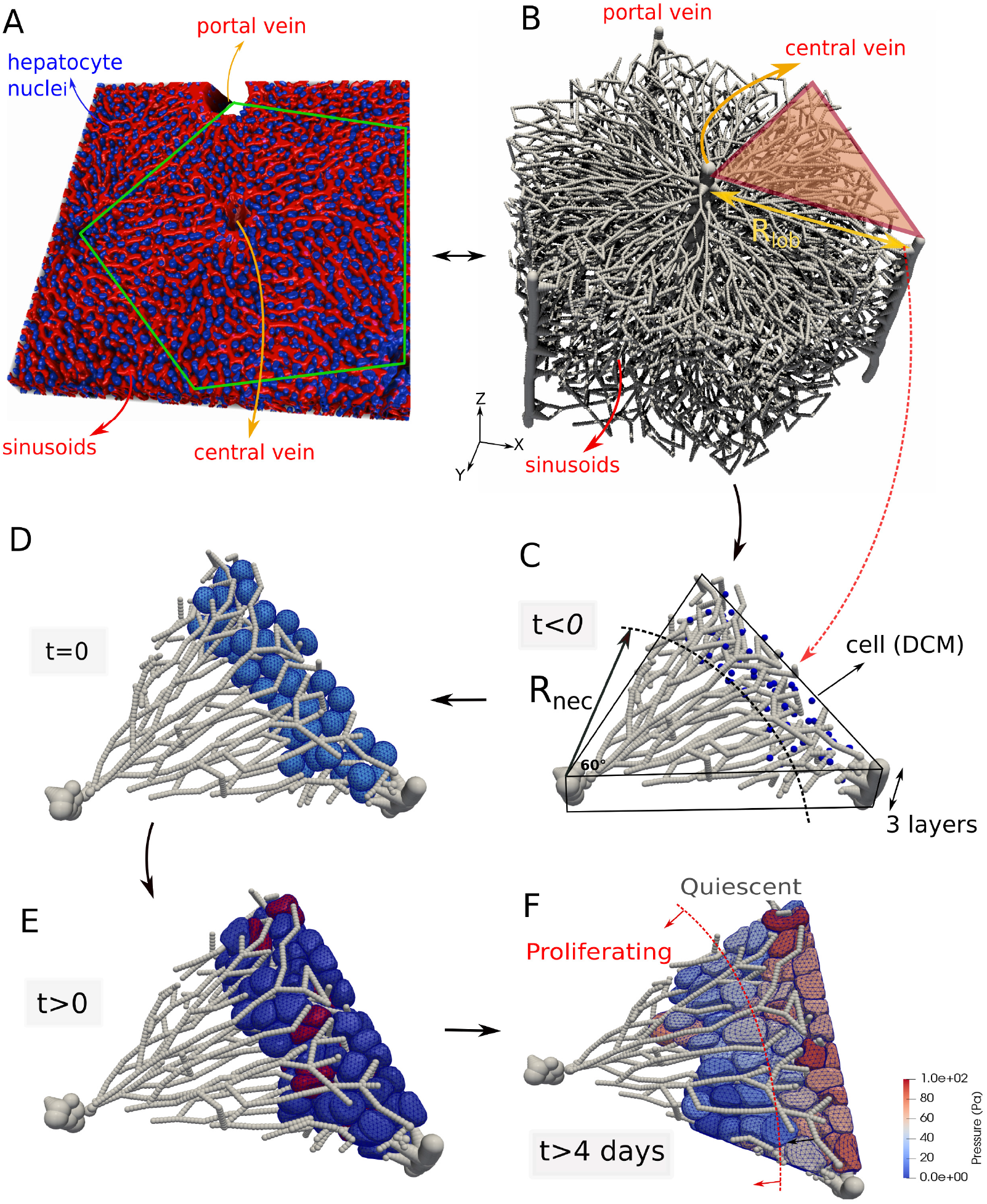
(A) Segmentation of hepatocyte nuclei (blue) and sinusoidal network (red) in liver tissue of mouse. Green lines outline a liver lobule sub-structure in the tissue. (B) Structure of the blood vessel network of a single lobule within the model. (C) Part of the network used for the simulations with the DCM. (C-D) Cells artificially growing up to realistic cell size (initialization). (E) The spatial-temporal proliferation pattern is imposed. The red cells are dividing. (F) Regeneration process the lobule, cells are colored according to internal pressure.

As in ref. [4] we used a three-cells-thick lobule architecture representative of a mouse liver. The sinusoids and veins are approximated as a dense network of fixed overlapping spheres, each with radius equal to the vessel radii. The high density of overlapping spheres warrants a smooth interaction with the cells. The hexagonal shape allows for identification of 6 statistically identical “pies” together constituting a full lobule. As a full lobule is currently not amenable to simulations with the DCM because of the large computation times, we simulate regeneration in only one pie. At the outer boundaries of the lobule pie, the cells are pushed back by vertical planes that mimic the presence of the neighboring lobule parts, see Fig. 8B. In our model, we assume that vessels can interact with the cells, yet their positions remain static during the simulation [4]. We perform both DCM and CBM simulations of the regeneration and quantitatively compare the differences. Both simulations start from identical initial conditions and configurations, and have the same model parameters (see Table 1). A maximum equivalence of mechanical parameters for both model types is ensured (see section 2.3).

We start from a multi-cellular configuration characteristic for the beginning of the regeneration process after CCl4 (or acetaminophen)-generated damage, where only hepatocytes localized at distance of at least *R_nec_* from the central vein survive (see Fig. 8C-D, blue colored cells). Hepatocytes localized at distances smaller than *R_nec_* from the central vein are assumed to be killed by the drug and removed from the system as those express the CYP-enzymes metabolizing CCl4 to NAPQI, which downstream leads to cell death. At *t <* 0, there are about 90 cells positioned in the free spaces between the sinusoids. An initial small radius is assigned to the hepatocytes, so that they initially do not touch the sinusoids. Next, the hepatocytes’ size is artificially increased, gradually generating contact with the sinusoids, until the hepatocytes reach a radius of *R_c_* = 12 *µ*m (Hammad et al., 2014) and the system a mechanical equilibrium (Fig. 8D). The time at this starting configuration is set to *t* = 0 as we here focus on the regeneration process (which is about 2-2.5 days after drug administration [4]).

During regeneration, those cells that have survived the intoxication enter the cell cycle to grow and divide with a rate depending on their distance from the necrotic lesion. During the regeneration process, they gradually move towards the central part, and eventually close the lesion and restore the lobule hepatocyte mass. Hepatocytes do not enter as individual, isolated cells (like mesenchymal cells) but in a collective movement as a sheet (with epithelial phenotype). In the simulations performed in [4] the spatio-temporal proliferation pattern of the cells during regeneration from an experimental *BrdU* staining pattern was directly imposed to the cells. *BrdU* stains cells in S-phase. Here, we impose a similar but a slightly coarse-grained pattern (for simplicity and better comparability to CBM-simulations), where cells are proliferating in a certain time window (Fig. 10A) and in a region smaller than 4-5 cell layers closest to the necrotic lesion. Outside this window, cells are assumed to be quiescent. Despite we do not simulate the full lobule, we nevertheless ensure that the ratio of the cells numbers before intoxication, after intoxication, and after regeneration are comparable as in ref. [4], so they would lead to the same numbers if extrapolated to an entire lobule.

The cells gradually progress towards the central vein to close the lesion with the largest fraction of proliferating cells at the border to the lesion (the front of the expanding tissue) and smallest proliferation activity at the periportal field (Fig. 8F). During the simulations, various cell state variables such as pressure, volume, state and tissue properties (cell density, area of the necrotic zone) can be monitored (see Fig. 8,Fig. 10 and Fig. 9D-E)

**Fig 9.**
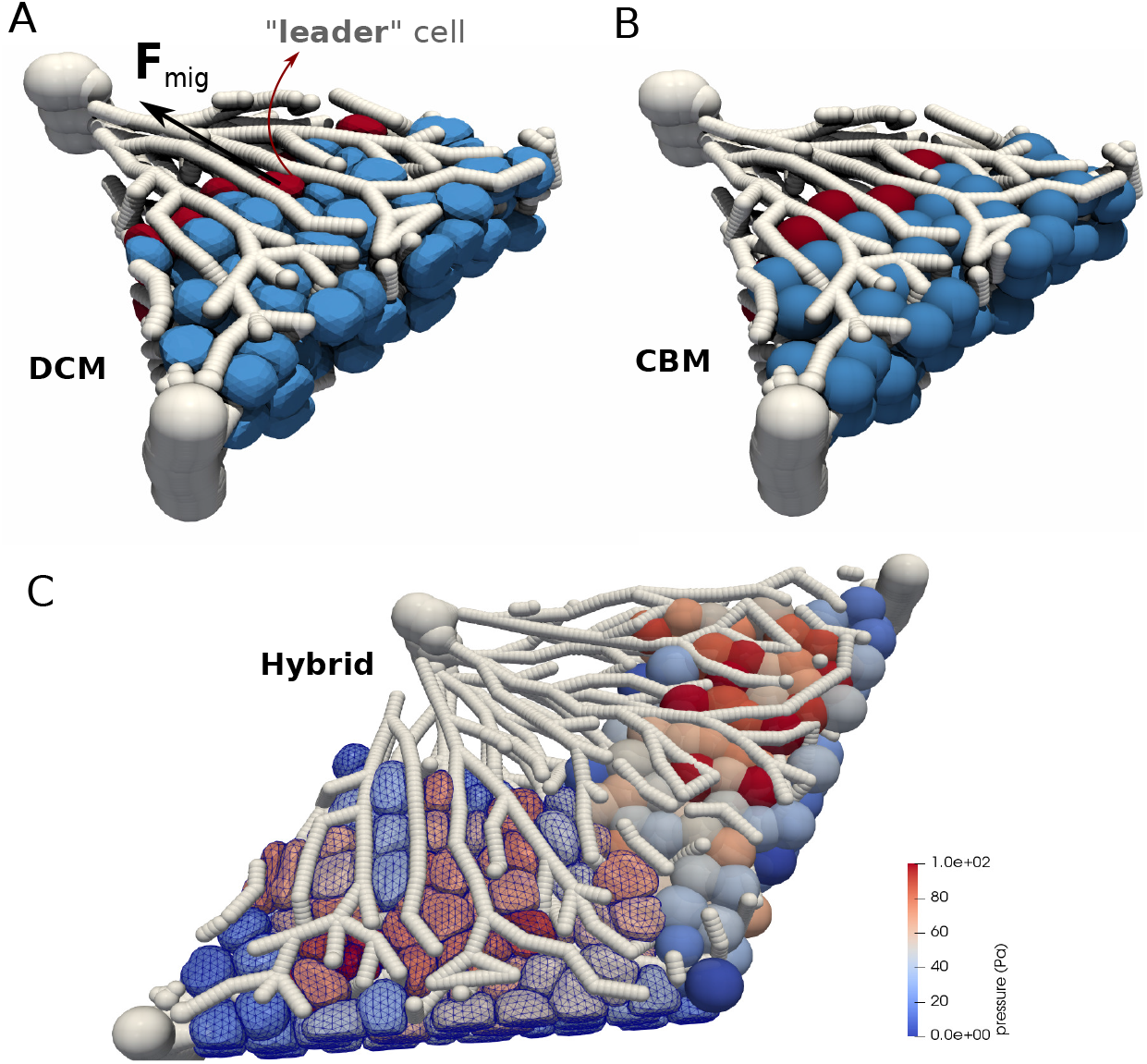
Simulation snapshots of the liver regeneration performed with the DCM (A) and CBM (B). (C) Simulation of CBM and DCM in hybrid mode. The colorbar here is according to cell pressure. Videos 3 and 4 of this in silico experiment are provided in the SI 6.3.

**Fig 10.**
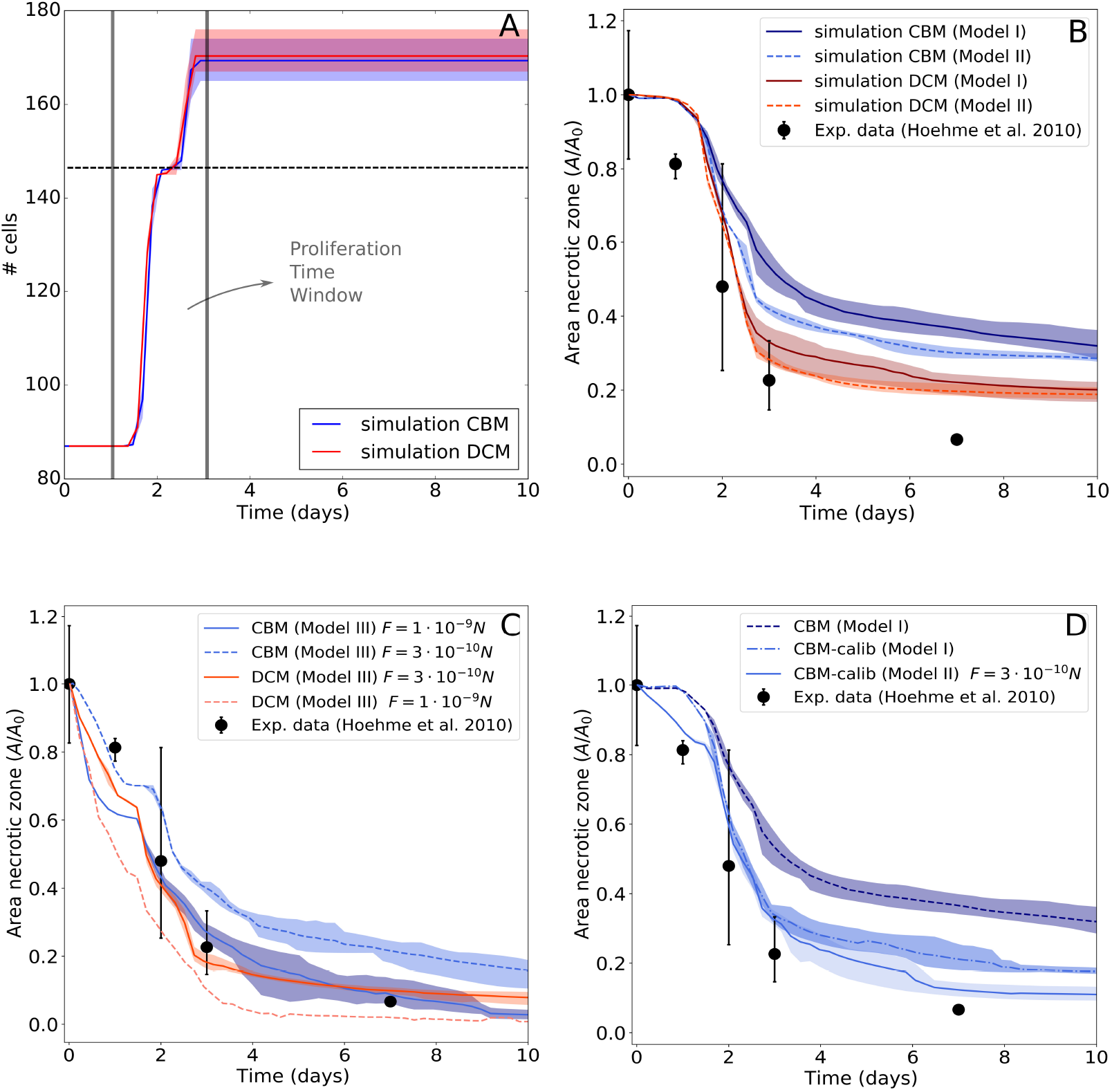
(A) Simulation (DCM and CBM) of the evolution of cell numbers during the time course of the simulation. (B) Simulated (DCM and CBM) relative area of the necrotic lesion during liver regeneration compared to data from Hoehme et al. (2010), assuming Model I or Model II. (C) Simulated (DCM and CBM) relative area of the necrotic lesion assuming Model III. (D) Simulated (CBM) relative area of the necrotic lesion using the calibration procedure for contact forces, assuming Model I or Model III. Each line is the average of 5 simulations with different random seeds, the shadowed zone indicates maximum and minimum values.

In a next step we study whether different hypotheses addressing the behavior of individual cells can explain the closure of the pericentral liver lobule lesion generated by toxic concentrations of CCl4 or acetaminophen. Guided by the choices in ref. [4] we distinguish between three model cases in which the cells differ by certain migratory or proliferation-related properties. In the basic model (Model I), we assume the cells to be proliferating with daughter cells oriented in a random direction and active movement generated by an unbiased random force (controlled by the cell diffusion coefficient *D*)^7^. We considered the lesion closure speed^8^ in Fig. 10B, which shows that this model is not in agreement with the experimental data for a time window of 10 days after intoxication, for both the DCM and CBM. This is in line with the conclusion in ref. [4], stating that the pericentral lobule lesion does not completely close within the experimentally observed time period if the driving force for the closure is cell proliferation only. Nevertheless, the DCM simulations exhibit a closer agreement with the experimental data than the CBM. Potentially, this is due to the more realistic contact forces and cell shapes in DCM compared to the CBM, which permit deformable cells to adapt their shape by squeezing in between the sinusoids. The pairwise JKR-based contact force in CBM lumps force contributions originating from cell deformation and compression into one mathematical formula assuming slight cell shape changes only. As a consequence at large compression forces emerging as a result of massive cell proliferation as in liver regeneration after drug-induced pericentral damage, significant overlap among cells, and cells and blood vessels can occur. The overlap might lead to an underestimation of the total volume occupied by the cells and blood vessels and too small repulsion forces between cells, and cells and vessels. This in turn critically slows down cell movement of cells that are pushed by proliferating cells towards the lesion, as the pushing forces result from volume compression. In the DCM simulations, volume is explicitly tracked hence cells are pushed away stronger. At the same time, DCM-cells can automatically adapt their shape and squeeze through the network. Both effects together speed up the cell displacement in DCM compared to CBM.

In the second model (Model II), we assumed that the during division the daughter cells align along the closest sinusoid, in Hoehme et al. (2010) [4] named “hepatocyte-sinusoid alignment” (HSA) mechanism and identified as responsible order mechanism for the reconstitution of liver lobule micro-architecture. In ref. [4] healthy liver lobule micro-architecture was shown to be characterized by a maximal hepatocyte-sinuoid contact area, facilitating the molecular exchange area between hepatocytes and blood. However, contrary to the model of Hoehme et al., where cell polarity was introduced as anisotropic adhesive forces, we here do not impose cell polarity; HSA in combination with differential adhesion between hepatocytes and lack of adhesion between hepatocytes and sinusoids already result in a largely columnar order. Moreover, in Hoehme et al. (2010) HSA was combined with a directed migration of hepatocytes towards the central necrotic lesion, which we drop in Model II to test the effect of HSA only. Interestingly, with this “directed division” mechanism, we observe for both the CBM and the DCM a slight “speed-up” of the lesion closure, in line with the reasoning that cells get less obstructed by the sinuoids after division if aligned. Yet, an acceptable quantitative agreement is not obtained Fig. 10B.

Finally, in the third model (Model III) we assume that cells migrate towards the central vein as a response on a morphogen gradient, whereby the morphogen source is the necrotic lesion. The directed migration manifests itself as an active force **F**^*mor*^ in the equation of motion (see section 2.1). Cell migration in liver is mediated by the extracellular matrix (ECM) scaffold, whereby the ECM is localized mainly in the space of Disse [78]. In Hoehme et al. (2010), hepatocytes at the border of the necrotic lesion were observed to stretch filopodia into the necrotic lesion. Consequently, the migration force **F**^*mor*^ is assumed to only act on the outer cells (here referred to as “leader” cells) and directed towards the central vein.

In Hoehme et al. [4] adding directed migration forces towards the central vein to Model I was able to ensure complete lesion closure.

For the DCM simulations with Model III, we found an excellent agreement with the data if the migration force had a strength of *F ^mor^* ∼ 0.3 nN. Contrary, for the CBM simulations with the same model (Model III) a significantly larger force of 1 nN is needed to close the lesion, showing again the significant and important differences between the two model approaches at the quantitative level (Fig. 10C). In turn, simulations with DCM with the migration force chosen as the migration force needed in the CBM to close the lesion (i.e. 1 nN) led to a too fast motion of the hepatocytes, hence a too fast closure of the lesion (Fig. 10C). This again confirms DCM-cells adapt easier to the obstructions imposed by the environment. However, overall the magnitude of the migration force for both approaches is in agreement with experimental observations of migrating cells [79, 80]. Specific measurements for regeneration after drug induced damage have so far not been performed.

The simplified formalism in CBMs thus lead to unadapted cell shapes (Fig. 9A-B) and possibly quantitatively incorrect contact forces and cell density. In order to test the influence of the contact forces between the cells on the results, we applied our procedure to re-calibrate the CBM contact forces from the DCM (see section 2.3.1 and section 6.1). In the procedure we simulate a spheroid compression experiment. The contact stiffness of every CBM cell is adjusted depending on by how many neighbor cells it is surrounded and by the relative distances between the cell and its neighboring cells, to finally obtain the same contact forces as in the equivalent DCM simulation. As such, we obtain a calibrated CBM that takes the effect of multiple cell contacts into account. Using the calibrated CBM, we rerun the simulations for Model I and Model II. For Model I, the lesion is now closing faster and comparable to the DCM simulations. For Model III we now obtained a much better agreement if using the same migration force as in DCM (Fig. 10D). This overall shows the significant impact of quantitatively correct contact forces for quantitative simulations of the cell dynamics in the liver lobule. Simulations with center-based models may still be used to give valuable quantitative insight if the cell contact mechanics is corrected. The DCM thus permits to partially correct for inaccuracies of the CBM, and to serve as an instrument for verification simulations for selective parameter sets, whereby parameter sensitivity analysis simulations might now still be performed with the computationally much faster CBM. We conclude that with a more refined cell model as the DCM model, the magnitude of the active migration force needed to close the central necrotic lesion after drug-induced peri-central damage by CCl4 is smaller than for the center-based model, but over all the same mechanisms i.e., active hepatocyte migration towards the necrotic lesion and oriented cell division are necessary for closure of the lesion. This qualitatively confirms the conclusion made in Hoehme et al. (2010) based on simulations with a CBM. However as we demonstrated, the differences between CBM and DCM simulations become very small if the cell-cell interaction forces in the CBM are calibrated by simulations with the DCM.

Finally and in line with this argument, we demonstrate that the two model approaches are capable to work in a hybrid mode. This is particularly useful if one can subdivide the total system in zones with a higher interest where more detail is required (high interest), and others where only approximated dynamics is needed (low interest), see e.g. [81]. Here, a lower computational cost can be obtained if the zone of lower interest can be simulated using CBM. Thanks to the nature of the contact model of the DCM, we can run simulations where parts of the lobule are covered by a DCM while others by a CBM. In Fig. 9C we considered two equal parts of the liver lobule where each part is covered by another model. At the interface, the model ensures that cells from DCM and CBM smoothly interact.

## 4 Summary and conclusions

In this paper our main goal was to study in how far model-based statements and interpretations on regeneration after drug induced liver damage depend on the degree of detail in the representation of hepatocyte shape and mechanics. For this purpose we developed a computational model that permits to explicitly mimic the cell shape and subcellular details.

We elaborated on a previously introduced deformable cell model (DCM) [17, 32] that has been proven to simulate realistic cell shapes in response to local internal and external forces. The *in silico* cells were constructed by a nodal network of viscoelastic elements that represent the cortical cytoskeleton and contain a homogeneous compressible cytoplasm. In this paper we have extended this model with cell proliferation and motility to perform multi-cellular simulations, with a model for cell growth and division. We demonstrated how the viscoelastic elements can be chosen to explain data at time scales differing by order of magnitudes, namely optical stretcher experiments to probe cell biomechanics on a time scale of seconds, as well as tissue regeneration occurring at time scales of several days.

An adequate choice of the rheological model for the individual subcellular elements (representing the outer cytoskeleton) and the calibration of parameters hereof, is an essential aspect for acquiring realistic simulations. Starting from simulating deformation of MDA-MB-231 cells in an optical stretcher experiment, we have shown that commonly used linear spring damper elements like Kelvin-Voigt and Maxwell may be sufficient to reproduce experimental data over a short time course. Overall, the Maxwell model approach seems to be more realistic in predicting cell relaxation behavior. In line with the analysis in [64], changes of friction coefficients in response to the ambient experiment temperature changes were necessary. Performing in silico pull-off experiments imposing increasing adhesion energies, we have further verified that the adopted cell contact model consistently reproduces realistic forces in cell-cell adhesive contacts. Here, the model shows a further potential for investigating local stress distributions in cell-cell adhesive bonds, which can be important to understand mechano-transduction processes.

Cell growth and division occurs on much longer time scales than those probed by most experimental methods (Optical Stretcher or Atomic force microscopy). Simulating cytokinesis explicitly by a contractile ring that splits the mother cell body into two approximately equal daughter cells, requires a re-meshing of the cell surface which turned out to be algorithm-wise very complex. Our final algorithm in a first step grows a cell to the twofold of its initial volume immediately after division keeping the number of nodes *N_s_* on the surface constant. In a second step for cell division, two cell bodies are inscribed in the original volume of the mother cell with *N_s_* surface nodes each, increasing their volume until the volume of the mother envelop cell is precisely filled by its two daughter cells, after which the latter is removed.

The ability in the model for cells to grow and divide allows to simulate small in-vitro experiments such as monolayers and spheroids growth up from a single cell up to 1000 cells on a single processor of a standard PC in about 1 week time. Despite these relatively low cell numbers, we have here illustrated that adhesion energy variations between cells and substrate may have different effects in monolayers as compared to spheroids.

Hoehme et al. (2010) demonstrated that agent-based models of the center-based type, mimicking cell-cell forces as forces between cell centers, are capable of giving valuable insights in liver regenerative processes. In particular, their simulations of a liver lobule regeneration showed that after intoxication, cell cycle re-entrance and cell division alone cannot explain the closure of lesions. By iterations between model simulations and experiments, they concluded that cells need migrative forces in order to acquire a fully regenerated lobule in a realistic time course. Our simulations with the more sophisticated cell model confirm this hypothesis, yet comparisons between our model and the center-based approach indicates some significant quantitative differences, which we could use to identify critical model determinants that must be represented in sufficient detail.

1. In the DCM, cells adapt to their environment requiring lower migrative forces to close the gap as compared to their center-based counter parts. In the latter, large cell-cell overlaps exists and rigid cell shapes result in a lower closure speed or higher required migrative force to close the lesion.
2. Cell contact forces play an important role in the dynamics. Replacing the (standard) JKR contact force model in the CBM by new force relations obtained from simulated force probing experiments with the DCM largely removed the difference in the regeneration dynamics between CBM and DCM.
3. DCM simulations result in more accurate cell shapes along the sinusoids as they adapt to the environment (cells do not overlap with sinuoids), and can in principle even directly be performed from confocal micrographs (see example in Fig. 11).

**Fig 11.**
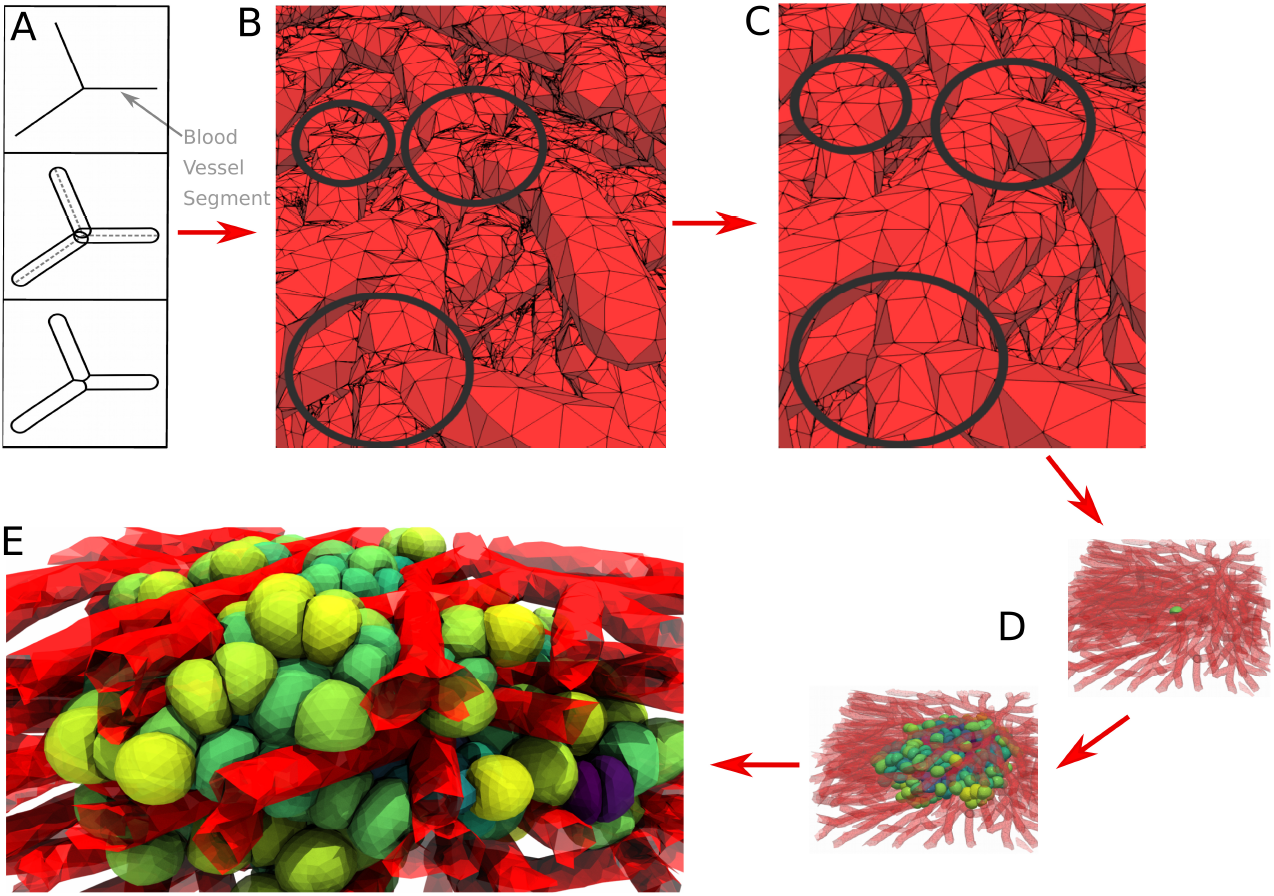
Simulation snapshots of a prototype model for liver tissue where the sinuoidal architecture and vessel shapes inferred from confocal micrographs. (A) We first approximate each blood vessel section as a cylinder with rounded ends. We rotate and translate them to match them with the position and direction of the original segment. The cylinders are then meshed. We use the meshing algorithms available in the C++ CGAL library [82] (B) In a next step (C) we compute with CGAl the intersection of all vessel meshes and remove artifacts (indicated by circles) due to the intersection algorithm. Finally, we obtain a realistically meshed bloodvessel network that allows interaction with a growing DCM cells (D-E).

In conclusion, we have shown that this highly detailed model approach may become a requirement to simulate tissue regeneration quantitatively, integrating subcellular information on cell shapes and biomechanics. For the future, we strive to run simulations directly starting from triangulated structures (representing cells, blood vessels, flexible or stiff structures, see prototype simulation in Fig. 11), which can be imported automatically from processed images [83] without further approximations or assumptions about shape. This is likely to increase the computational cost and require distributed computing. Large population DCM simulations will require code parallelization envisaged for the future. Yet, the approach proposed here also allows to simulate tissues in a hybrid mode. This may open possibilities to accomplish large scale tissue simulations where some parts are modeled at high spatial resolution models, while in other zones approximate models are sufficient. We have demonstrated this by hybrid simulations in a proof of concept.

## 5 Acknowledgments

PvL, JN, TJ implemented code, PvL and JN performed model simulations, PvL, JN and DD developed the model, PvL, JN and DD: analyzed the simulation data. PvL SH, JK and DD guided research, EW and SG performed experiments, EW, SG, JK analyzed experimental data, PvL and DD wrote the paper, DD guided the project.

PvL and DD would like to thank Manuel Garcia Aznar for his valuable input.

PvL and DD gratefully acknowledges support by BMBF - LiSym (DD), INST. CANCER - PHYSCANCER (DD), ITMO - INVADE (DD), EU 7th Framework Programme (NOTOX), and ANR - iLite. JN and TJ gratefully acknowledge support by BMBF-LiSym (DD), BMBF-VLN (DD) and BMBF-Demonstrator Liversimulator (DD) SH gratefully acknowledges support by DFG-Emmy-Noether and BMBF-VLN (DD). EW, SG and JK … DD furthermore gratefully acknowledges support by BMBF VLN and Demonstrator Liversimulator. JS, EW and SG greatly acknowledge the support of the ERC grant nr 741350.

## 6 Supplementary Material

### 6.1 Determining DCM contact forces and friction coefficients

A small cell clump was used as *in silico* model to estimate the contact forces and relaxation in a cell aggregate. We considered a central cell neighbored by several others initially structured as a body centered cubic (Fig. 12A). The cell clump was then positioned in a imaginary rigid spherical capsule. The radius of the capsule was steadily and step-wise decreased, compressing the cells inside (Fig. 12A). The average distance between the central cell and its contacting cells was computed as *d_ij_* = 1 *d_ij_ /*(*R_i_* + *R_j_*) at every timestep. This resulted in a forces-distance relation during the time course of compression that was used to calibrate the contact forces in the center-based model. In Fig. 12B we have displayed the average force-distance relation during the experiment. The contact force in the center-based model shows a clear deviation as the distance gets smaller. This clearly shows that JKR or Hertz contact force models do not apply for high cell densities in CBM.

**Fig 12.**
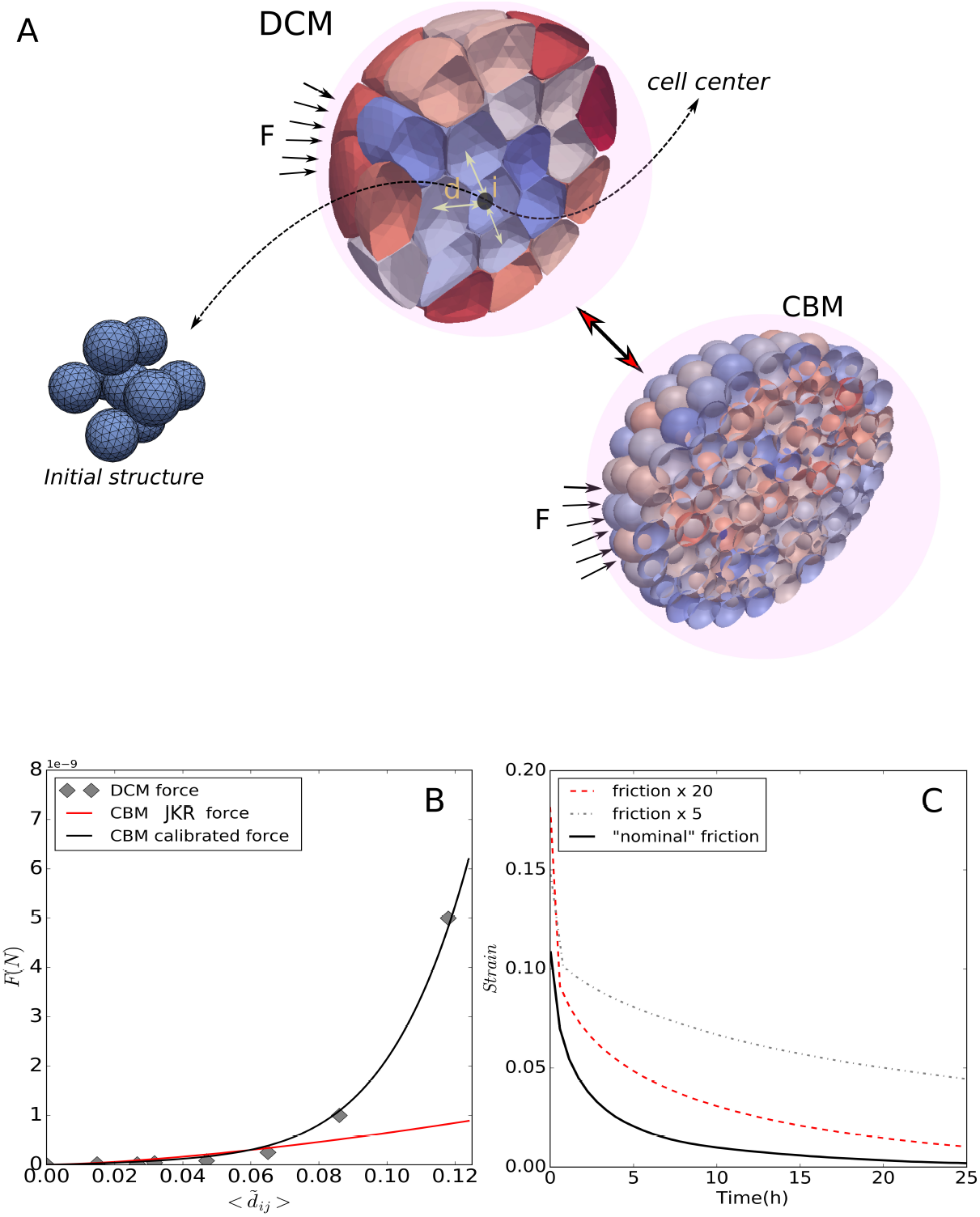
(A) Snapshot of a simulation of a small cell clump of DCM being compressed by isotropic forces. The distance *d* are computed between the cell centers and averaged. The equivalent CBM simulation is depicted below. (B) Force-distance data for a compression experiment simulated with DCMs, CBMs with using the JKR contact model, and CBMs using the calibrated contact model. (C) Simulated relaxation curves of a small spheroid performed with DCMs assuming different friction coefficients. Video 2 of this *in silico* experiment is provided in the SI 6.3.

The full relaxation behavior of the spheroid is determined by the nodal friction coefficients, the viscous friction of the adherent cells, and the friction with the ECM. To measure relaxation, the cells are suddenly released from the encapsulation. This period for the cells to come back to their original state is a measure for the relaxation time of a spheroid, and is used to calibrate the friction coefficients. The relaxation state is quantified by a strain function measuring again the average distance function *d* between the cells as a function of time. This is depicted in Fig. 12C. We choose the optimal friction coefficients so that the cell clump has a relaxation time of ∼ 5*h* as mentioned in [75, 76]. A model run with 5 times higher and 20 times friction is depicted to show a significant increase of relaxation times.

### 6.2 Estimation of the surface forces on a cell during the optical stretcher experiment

Here, we give explain how the nodal forces on the cell are calculated from the optical laser beam. Therefore, we closely follow the approach detailed in [36]. The cells pass the optical stretcher experimental setup in suspension. The radius of the Gaussian laser beam is *R_beam_* = 8.2 *µ*m [63]. We can assume a ray optics approach, since the wavelength of the laser light, *λ* = 1064 nm is much smaller than the diameter of our optical particles (≈ 17*µm*).

Assume first a cube-like object which obstructs in the laser beam path. The laser beam enters the cube at the front side and leaves it at the back side. The incident momentum of the laser *p_i_* = *nE/c* (E is the energy of the incoming beam, *n* is the refractive index and *c* is the speed of light)), needs to be conserved at the interfaces of the medium with the cube. The surface picks up the difference of momentum ∆*p* = *p_incident_* + *p_ref lected_ p_transmitted_* between the incident ray and the transmitted ray through the cube.

A resulting optical force acting due to momentum transfer for the frontal cube side is calculated using the Fresnel formulas and Newton’s second law [36]:

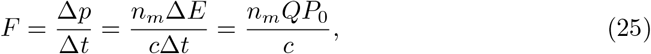

where *E* is the energy of the photons, *c* the speed of light, *n_m_* is the refractive index of the medium. The factor *Q* represents the transferred momentum (*Q* = 2 for total reflection, *Q* = 1 for total absorption) and total *P*_0_ is the incident power. The same formula needs to be applied to the laser beam leaving the cube to the medium. The total force on one side of the cube at the laser axis due to the presence of two lasers in opposite direction can thus be calculated by summing up the transferred momenta at one side [36, 84]:

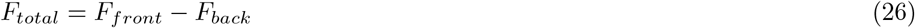

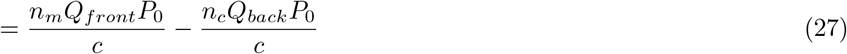

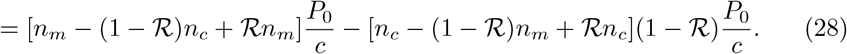

where *n_c_* is the refractive index of the cube. The reflection can be neglected for small incident angles, since according to the Fresnel Equation:

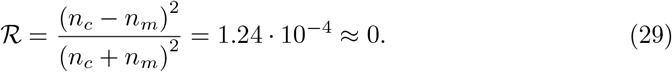

Hence the total force on the cube interface simplifies to:

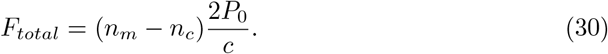

This force acts always perpendicular to the object opposed to the incoming laser light. The applied stress in the direction of the laser beam on the cube is thus:

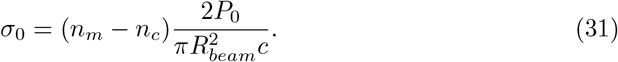

For *n_m_* = 1.33, *n_c_* = 1.36, *P*_0_ = 1.1W and *c* = 3 10^8^ m*/*s, we find *σ*_0_ = 1.05 Pa.

Within our model we need the applied stress for each triangular surface segment. However, the forces due to the laser beam are not equally distributed over the cell, as the cell is not a cube but has a spherical-like shape and the incident angle is not zero anymore but varies along the cell surface. A realistic overall stress profile can be assumed to be approximated by *σ*(*α*) = *σ*_0_*cos^n^α* with amplitude *σ*_0_ and fitting parameter *n* [68]. Here, *α* is the angle between the laser axes and the nodal position of the cell (see Fig. 4C). To determined the parameter *n*, we evaluate the ratio of the laser beam radius and cell radius:

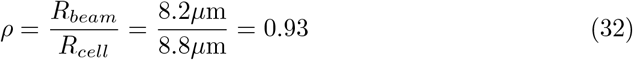

For a ratio of *ρ ≈* 1, *n* fits best when set to 2 (see e.g. [36, 68, 84]). As we know how much surface area *A_n_* each node on the cell represents, we can now calculate the forces on the DCM nodes by *F_L_*(*α*) = *A_n_σ*(*α*), where *F_L_*(*α*) is oriented perpendicular to the cell surface at that node (Fig. 4C).

### 6.3 Videos

**Video 1.** Simulation of pull-off experiment with DCM. The cells move in opposite directions due to opposing forces that eventually break the adhesive bond.

**Video 2.** Simulation of monolayer growth with DCM starting from 1 cell.

**Video 3.** Simulation of liver regeneration using DCM with a migrative force of 3 nN.

**Video 4.** Simulation of liver regeneration using CBM with a migrative force of 10 nN.

**Video 5.** Simulation the compression experiment with DCM to determine the contact forces between the cells and the influence of the friction coefficients on the relaxation.

## Footnotes

1 In this paper we assume that friction with ECM is isotropic and variations in cell shape do not alter the friction significantly.

2 We assume here a constant and homogeneous adhesion field. However experiments have pointed out that adhesion bounds are more point-like and can re-inforce over time [50]. Although this considerations could be addressed with our model, it remains out of the scope of this paper.

3 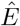 is given a large value compared to the cell Young modulus to prevent interpenetration of triangles

4 A sub-simulation means that it runs in parallel and does not add up to the total simulation time (time is kept constant). The duration of a sub-simulation is very short, of the order of 10s.

5 Because of the presence of bending elasticity and resistance against area expansion of the cell plasma membrane, the overall behavior is not purely fluid-like but may rather be regarded as a Standard Linear Solid (SLS). However, these effects only appear on much longer timescales in the simulations.

6 The long-term relaxation into a spherical shape might be mimicked by implementing an additional spring with small spring constant in the MME element.

7 The assumptions in Model I are equivalent to those in Model 1 in ref. [4]

8 We define here the lesion radius as the average distance from the central vein to the closest cell in every layer whereby we consider the 3 closest layers to the lesion.

## References

1. Fletcher DA, Mullins RD. Cell mechanics and the cytoskeleton. Nature. 2010;463(7280):485.

2. Rodriguez ML, McGarry PJ, Sniadecki NJ. Review on cell mechanics: experimental and modeling approaches. Applied Mechanics Reviews. 2013;65(6):060801.

3. Karolak A, Markov DA, McCawley LJ, Rejniak KA. Towards personalized computational oncology: from spatial models of tumour spheroids, to organoids, to tissues. Journal of The Royal Society Interface. 2018;15(138):20170703.

4. Hoehme S, Brulport M, Bauer A, Bedawy E, Schormann W, Hermes M, et al. Prediction and validation of cell alignment along microvessels as order principle to restore tissue architecture in liver regeneration. Proceedings of the National Academy of Sciences. 2010;107(23):10371–10376.

5. Drasdo D, Höhme S. A single-cell-based model of tumor growth in vitro: monolayers and spheroids. Physical biology. 2005;2(3):133.

6. Drasdo D, Hoehme S, Block M. On the Role of Physics in the Growth and Pattern Formation of Multi-Cellular Systems: What can we Learn from Individual-Cell Based Models. Journal of Statistical Physics. 2007;128:287–345.

7. Geris L, Van Liedekerke P, Smeets B, Tijskens E, Ramon H. A cell based modelling framework for skeletal tissue engineering applications. Journal of biomechanics. 2010;43(5):887–892.

8. Buske P, Galle J, Barker N, Aust G, Clevers H, Loeffler M. A Comprehensive Model of the Spatio-Temporal Stem Cell and Tissue Organisation in the Intestinal Crypt. PLoS Comput Biol. 2011;7(1):e1001045.

9. Ramis-Conde I, Chaplain MAJ, Anderson ARA, Drasdo D. Multi-scale modelling of cancer cell intravasation: the role of cadherins in metastasis. Physical Biology. 2009;6:16008.

10. Delile J, Herrmann M, Peyriéras N, Doursat R. A cell-based computational model of early embryogenesis coupling mechanical behaviour and gene regulation. Nature communications. 2017;8:13929.

11. Drasdo D, Hoehme S, Hengstler JG. How predictive quantitative modelling of tissue organisation can inform liver disease pathogenesis. Journal of hepatology. 2014;61(4):951–956.

12. Schaller G, Meyer-Hermann M. Multicellular tumor spheroid in an off-lattice Voronoi-Delaunay cell model. Phys Rev E. 2005;71(5):51910.

13. Van Liedekerke P, Palm MM, Jagiella N, Drasdo D. Simulating tissue mechanics with agent-based models: concepts, perspectives and some novel results. Comp Part Mech. 2015;2(4):401–444.

14. Rejniak Ka. An immersed boundary framework for modelling the growth of individual cells: an application to the early tumour development. Journal of Theoretical Biology. 2007;247(1):186–204.

15. Sandersius SA, Newman TJ. Modeling cell rheology with the Subcellular Element Model. Physical biology. 2008;5(1):15002.

16. Jamali Y, Azimi M, Mofrad MRK. A sub-cellular viscoelastic model for cell population mechanics. PLoS One. 2010;5(8).

17. Odenthal T, Smeets B, Van Liedekerke P, Tijskens E, Van Oosterwyck H, Ramon H. Analysis of initial cell spreading using mechanistic contact formulations for a deformable cell model. PLoS computational biology. 2013;9(10):e1003267.

18. Graner F, Glazier JA. Simulation of Biological Cell Sorting Using a Two-Dimensional Extended Potts Model. Physical Review Letters. 1992;69:2013–2016.

19. van Oers RFM, Rens EG, LaValley DJ, Reinhart-King Ca, Merks RMH. Mechanical Cell-Matrix Feedback Explains Pairwise and Collective Endothelial Cell Behavior In Vitro. PLoS Comput Biol. 2014;10(8):e1003774.

20. Palm MM, Merks RMH. Vascular networks due to dynamically arrested crystalline ordering of elongated cells. Phys Rev E. 2013;87(1):12725.

21. Dillon R, Owen M, Painter K. A single-cell-based model of multicellular growth using the immersed boundary method. AMS Contemp Math. 2008;466:1–15.

22. Rejniak KA, Anderson ARA. A computational study of the development of epithelial acini: I. Sufficient conditions for the formation of a hollow structure. Bulletin of Mathematical Biology. 2008;70(3):677–712.

23. Hosseini M, Feng JJ. A particle-based model for the transport of erythrocytes in capillaries. Chemical engineering science. 2009;64:4488–4497.

24. Fedosov DA, Caswell B, Karniadakis GE. Systematic coarse-graining of spectrin-level red blood cell models. Computer Methods in Applied Mechanics and Engineering. 2010;199(29–32):1937–1948.

25. Van Liedekerke P, Smeets B, Odenthal T, Tijskens E, Ramon H. Solving microscopic flow problems using Stokes equations in SPH. Computer Physics Communications. 2013;184(7):1686–1696.

26. Fedosov Da, Caswell B, Suresh S, Karniadakis GE. Quantifying the biophysical characteristics of Plasmodium-falciparum-parasitized red blood cells in microcirculation. Proceedings of the National Academy of Sciences of the United States of America. 2011;108(1):35–9.

27. Sandersius SA, Weijer CJ, Newman TJ. Emergent cell and tissue dynamics from subcellular modeling of active biomechanical processes. Physical Biology. 2011;8:45007.

28. Van Liedekerke P, Tijskens E, Ramon H, Ghysels P, Samaey G, Roose D. Particle-based model to simulate the micromechanics of biological cells. Physical Review E. 2010;81(6, Part 1):61906–61915.

29. Van Liedekerke P, Roose D, Ramon H, Ghysels P, Tijskens E, Samaey G. Mechanisms of soft cellular tissue bruising. A particle based simulation approach. Soft Matter. 2011;7(7):3580.

30. Chen J, Weihs D, Van Dijk M, Vermolen FJ. A phenomenological model for cell and nucleus deformation during cancer metastasis. Biomechanics and modeling in mechanobiology. 2018; p. 1–22.

31. Fletcher AG, Osterfield M, Baker RE, Shvartsman SY. Vertex models of epithelial morphogenesis. Biophysical Journal. 2014;106(11):2291–2304.

32. Guyot Y, Smeets B, Odenthal T, Subramani R, Luyten FP, Ramon H, et al. Immersed boundary models for quantifying flow-induced mechanical stimuli on stem cells seeded on 3D scaffolds in perfusion bioreactors. PLoS Computational Biology. 2016;12(9):e1005108.

33. Tanaka S, Sichau D, Iber D. LBIBCell: a cell-based simulation environment for morphogenetic problems. Bioinformatics. 2015;31(14):2340–2347.

34. Milde F, Tauriello G, Haberkern H, Koumoutsakos P. SEM++: A particle model of cellular growth, signaling and migration. Computational Particle Mechanics. 2014;.

35. Turlier H, Audoly B, Prost J, Joanny JF. Furrow constriction in animal cell cytokinesis. Biophysical journal. 2014;106(1):114–23.

36. Guck J, Ananthakrishnan R, Mahmood H, Moon TJ, Cunningham CC, Kaes J. The Optical Stretcher: A Novel Laser Tool to Micromanipulate Cells. Biophysical Journal. 2001;81:767–784.

37. González-Valverde I, García-Aznar JM. Mechanical modeling of collective cell migration: An agent-based and continuum material approach. Computer Methods in Applied Mechanics and Engineering. 2018;337:246–262.

38. Kim MC, Neal DM, Kamm RD, Asada HH. Dynamic modeling of cell migration and spreading behaviors on fibronectin coated planar substrates and micropatterned geometries. PLOS Computational Biology. 2013;9(2):e1002926.

39. Tainaka K, Kuno A, Kubota SI, Murakami T, Ueda HR. Chemical principles in tissue clearing and staining protocols for whole-body cell profiling. Annual review of cell and developmental biology. 2016;32:713–741.

40. Sack I, Jöhrens K, Würfel J, Braun J. Structure-sensitive elastography: on the viscoelastic powerlaw behavior of in vivo human tissue in health and disease. Soft Matter. 2013;9(24):5672–5680.

41. Odell GM, Oster G, Alberch P, Burnside B. The mechanical basis of morphogenesis. I. Epithelial folding and invagination. Developmental biology. 1981;85(2):446–462.

42. Monnier S, Delarue M, Brunel B, Dolega ME, Delon A, Cappello G. Effect of an osmotic stress on multicellular aggregates. Methods. 2016;94:114–119.

43. Boal D. Mechanics of the Cell. 2nd ed. Cambridge University Press; 2012.

44. Sinha B, Köster D, Ruez R, Gonnord P, Bastiani M, Abankwa D, et al. Cells respond to mechanical stress by rapid disassembly of caveolae. Cell. 2011;144(3):402–413.

45. Van Liedekerke P, Buttenschoen A, Drasdo D. Off-Lattice Agent-Based Models for Cell and Tumor Growth: Numerical Methods, Implementation, and Applications. In: Numerical Methods and Advanced Simulation in Biomechanics and Biological Processes. Miguel Cerrolaza Sandra Shefelbine Diego Garzón-Alvarado; 2017.

46. Tinevez JY, Schulze U, Salbreux G, Roensch J, Joanny JF, Paluch E. Role of cortical tension in bleb growth. Proceedings of the National Academy of Sciences of the United States of America. 2009;106(44):18581–6.

47. Tamura K, Komura S, Kato T. Adhesion induced buckling of spherical shells. Journal of Physics: Condensed Matter. 2004;16(39):L421–L428.

48. Smeets B, Odenthal T, Keresztes J, Vanmaercke S, Van Liedekerke P, Tijskens E, et al. Modeling contact interactions between triangulated rounded bodies for the discrete element method. Computer Methods in Applied Mechanics and Engineering. 2014;277:219–238.

49. Maugis D. Adhesion of spheres: The JKR-DMT transition using a Dugdale model. J Colloid Interface Sci. 1992;150(1):243–269.

50. Pawlizak S, Fritsch AW, Grosser S, Ahrens D, Thalheim T, Riedel S, et al. Testing the differential adhesion hypothesis across the epithelial-mesenchymal transition. New Journal of Physics. 2015;17(8):083049.

51. Drasdo D, Forgacs G. Modeling the interplay of generic and genetic mechanisms in cleavage, blastulation, and gastrulation. Developmental Dynamics. 2000;219(2):182–191.

52. Johnson KL, Greenwood JA. An Adhesion Map for the Contact of Elastic Spheres. Journal of colloid and interface science. 1997;192:326–333.

53. Leckband D, Israelachvili J. Intermolecular forces in biology. Quarterly Reviews of Biophysics. 2001;34(2):105–267.

54. Tozluoğlu M, Tournier AL, Jenkins RP, Hooper S, Bates PA, Sahai E. Matrix geometry determines optimal cancer cell migration strategy and modulates response to interventions. Nature Cell Biology. 2013;15(7):751–762.

55. Kim MC, Silberberg YR, Abeyaratne R, Kamm RD, Asada HH. Computational modeling of three-dimensional ECM-rigidity sensing to guide directed cell migration. Proceedings of the National Academy of Sciences. 2018;115(3):E390–E399.

56. Drasdo D, Hoehme S. Modeling the impact of granular embedding media, and pulling versus pushing cells on growing cell clones. New Journal of Physics. 2012;14(5):55025.

57. Hammad S, Hoehme S, Friebel A, von Recklinghausen I, Othman A, Begher-Tibbe B, et al. Protocols for staining of bile canalicular and sinusoidal networks of human, mouse and pig livers, three-dimensional reconstruction and quantification of tissue microarchitecture by image processing and analysis. Archives of toxicology. 2014;88(5):1161–83.

58. Chu YS, Dufour S, Thiery JP, Perez E, Pincet F. Johnson-Kendall-Roberts Theory Applied to Living Cells. Physical Review Letters. 2005;94(2):28102.

59. Pathmanathan P, Cooper J, Fletcher A, Mirams G, Montahan L, Murray P, et al. A computational study of discrete mechanical tissue models. Physical Biology. 2009;6(3):36001.

60. Galle J, Loeffler M, Drasdo D. Modeling the effect of deregulated proliferation and apoptosis on the growth dynamics of epithelial cell populations in vitro. Biophysical journal. 2005;88(1):62–75.

61. Bock M, Tyagi AK, Kreft JU, Alt W. Generalized voronoi tessellation as a model of two-dimensional cell tissue dynamics. Bulletin of mathematical biology. 2010;72(7):1696–1731.

62. Guck J, Schinkinger S, Lincoln B, Wottawah F, Ebert S, Romeyke M, et al. Optical deformability as an inherent cell marker for testing malignant transformation and metastatic competence. Biophysical journal. 2005;88(5):3689–3698.

63. Grosser S, Fritsch AW, Kießling TR, Stange R, Käs JA. The lensing effect of trapped particles in a dual-beam optical trap. Optics Express. 2015;23(4):5221–5235.

64. Kießling TR, Stange R, Käs JA, Fritsch AW. Thermorheology of living cells—impact of temperature variations on cell mechanics. New Journal of Physics. 2013;15:45026.

65. Brugués J, Maugis B, Casademunt J, Nassoy P, Amblard F, Sens P. Dynamical organization of the cytoskeletal cortex probed by micropipette aspiration. Proceedings of the National Academy of Sciences of the United States of America. 2010;107(35):15415–15420.

66. Delarue M, Montel F, Vignjevic D, Prost J, Joanny JF, Cappello G. Compressive Stress Inhibits Proliferation in Tumor Spheroids through a Volume Limitation. Biophysical Journal. 2014;107(8):1821–1828.

67. Delarue M, Joanny JF, Jülicher F, Prost J. Stress distributions and cell flows in a growing cell aggregate. Interface focus. 2014;4(6):20140033.

68. Ananthakrishnan R, Guck J, Wottawah F, Schinkinger S, Lincoln B, Romeyke M, et al. Quantifying the contribution of actin networks to the elastic strength of fibroblasts. Journal of Theoretical Biology. 2006;242(2):502–516.

69. Kubitschke H, Schnauss J, Nnetu KD, Warmt E, Stange R, Kaes J. Actin and microtubule networks contribute differently to cell response for small and large strains. New Journal of Physics. 2017;19(9):093003.

70. Gyger M, Stange R, Kießling TR, Fritsch A, Kostelnik KB, Beck-Sickinger AG, et al. Active contractions in single suspended epithelial cells. European Biophysics Journal. 2014;43(1):11–23.

71. Murrell MP, Voituriez R, Joanny JF, Nassoy P, Sykes C, Gardel ML. Liposome adhesion generates traction stress. Nature Physics. 2014;10(2):163.

72. Schwarz US, Safran SA. Physics of adherent cells. Rev Mod Phys. 2013;85(3):1327–1381.

73. Sutherland RM. Cell and environment interactions in tumor microregions: the multicell spheroid model. Science (New York, NY). 1988;240(4849):177–84.

74. Brú A, Albertos S, Luis Subiza J, García-Asenjo JL, Brú I. The universal dynamics of tumor growth. Biophysical journal. 2003;85(5):2948–2961. doi:10.1016/S0006-3495(03)74715-8.

75. Marmottant P, Mgharbel A, Käfer J, Audren B, Rieu JP, Vial JC, et al. The role of fluctuations and stress on the effective viscosity of cell aggregates. Proceedings of the National Academy of Sciences of the United States of America. 2009;106(41):17271–5.

76. The Cellular Capsules technology And its applications to investigate model tumor. UPMC; 2013.

77. Byrne H, Drasdo D. Individual-based and continuum models of growing cell populations: a comparison. Mathematical Biology. 2009;58:657–680.

78. Mazza G, Rombouts K, Hall AR, Urbani L, Luong TV, Al-Akkad W, et al. Decellularized human liver as a natural 3D-scaffold for liver bioengineering and transplantation. Scientific reports. 2015;5:13079.

79. Kim MC, Whisler J, Silberberg YR, Kamm RD, Asada HH. Cell Invasion Dynamics into a Three Dimensional Extracellular Matrix Fibre Network. PLoS computational biology. 2015;11(10):e1004535.

80. Jacquemet G, Hamidi H, Ivaska J. Filopodia in cell adhesion, 3D migration and cancer cell invasion. Current opinion in cell biology. 2015;36:23–31.

81. Kim Y, Stolarska MA, Othemer HG. A Hybrid model for tumor Spheroid grwoth in vitro I: Theoretical development and earliy results. Mathematical Models and Methods in Applied Sciences. 2007;17(supp01):1773–1798.

82. The CGAL Project. CGAL User and Reference Manual. 4.13 ed. CGAL Editorial Board; 2018. Available from: https://doc.cgal.org/4.13/Manual/packages.html.

83. Friebel A, Neitsch J, Johann T, Hammad S, Hengstler JG, Drasdo D, et al. TiQuant: software for tissue analysis, quantification and surface reconstruction. Bioinformatics. 2015;31(19):3234–3236.

84. Guck J, Ananthakrishnan R, Moon T, Cunningham C, Käs J. Optical deformability of soft biological dielectrics. Physical review letters. 2000;84(23):5451.

